# Strong correlation between amino acid frequency and codon degeneracy in genetic codes across all domains of life

**DOI:** 10.64898/2026.09.10.750109

**Authors:** Valentin Wesp, Günter Theißen, Stefan Schuster

## Abstract

Since the discovery of the genetic code, a frequently discussed question is whether the numbers of synonymous codons for the various amino acids are randomly distributed or were shaped by evolutionary constraints. In this study, we analyze for the standard as well as alternative genetic codes, the correlations between codon degeneracy and amino acid frequencies in proteins (neglecting differences in gene expression). To taking into account the effect of GC content, expected codon frequencies rather than codon multiplicity need to be considered. A strong correlation of these frequencies with amino acid abundance is revealed. Furthermore, we identify consistent patterns of over- and underrepresentation of amino acids across domains of cellular life as well as viruses. For example, the codons for glutamate, aspartate, lysine, and methionine are consistently overrepresented across domains, while cysteine, arginine, histidine, and proline are underrepresented. Subsequently, we discuss the role of biosynthesis costs of amino acids and other factors such as the order of amino acid recruitment in early evolution, exposure to oxidative stress, and sulphur availability. We hypothesize that in a first phase of evolution, the genetic code evolved in a way so as to comply with the different demands for amino acids. In a second phase, after the code was frozen, changes in demand led to deviations from the strong correlation between codon multiplicity and amino acid frequency by natural selection. This comprehensive analysis offers new insights into the interplay between genetic code structure and amino acid usage across all domains of life.

## Introduction

The genetic code, a near-universal system across all biological species, maps nucleotide triplets (codons) to amino acids for protein synthesis. The code exhibits notable complexity: amino acids are encoded by varying numbers of codons, from a single codon for methionine and tryptophan each to six codons for leucine, arginine and serine each [1–3]. This disparity raises a fundamental question: Is there a consistently strong positive correlation between the number of synonymous codons for an amino acid and its overall genomic frequency? For example, do all leucine codons together appear six times as often in protein-coding genes as the codon for tryptophan? Or, once the code was fixed, did selection pressure shift amino acid frequencies away from that baseline distribution?

Amino acid frequencies in protein-coding genes across diverse species have been determined earlier (neglecting differences in gene expression) [4,5] and have been analyzed in relation to codon degeneracy [6–8]. Those studies revealed a noticeable positive correlation. Here, we start from the (even stricter) working hypothesis saying that these two quantities are proportional to each other.

Overall, an extensive body of scientific literature has explored the evolution of the standard and non-standard genetic codes [3,9–15]. A widely accepted theory posits that the extant genetic codes evolved in an early phase of evolution approximately three to four billion years ago (when the first life forms emerged), including the prebiotic era as well as an early biotic phase [10,15,16]. Codon degeneracy may have adapted to various constraints, notably the demand for the particular amino acids, their metabolic costs, availability and order of recruitment [16–19]. Throughout this paper, the terms “number of synonymous codons”, “codon degeneracy”, and “codon multiplicity” are used synonymously.

Simple amino acids, such as glycine, alanine, valine, serine, and leucine, are likely to Simple amino acids, such as glycine, alanine, valine, serine, and leucine, likely originated through abiotic synthesis, either in primitive environments [20] or via early RNA-based catalytic systems [15,16,18,19,21]. Due to their early availability, they were likely among the first incorporated into the genetic code and were assigned high numbers of codons [4,5,7,17,18,22]. In contrast, amino acids represented by a low number of synonymous codons, including methionine and cysteine (containing sulfur) as well as tryptophan (featuring an indole ring system) appear to have been incorporated later, when the genetic code was constrained to some extent already. This pattern correlates with the frequency of amino acids: simpler amino acids are more prevalent in proteomes, whereas structurally more complex amino acids occur at lower abundances [4,5,7].

Structural and functional requirements also play a significant role: hydrophobic residues are enriched in the core of cytosolic proteins as well as in membrane proteins, which affects amino acid abundance as well [7,23]. Additionally, a conserved bimodal distribution of isoelectric points across proteomes has been found and highlights evolutionary pressures balancing structural stability and functional flexibility [24]. Moreover, genomic properties, such as GC content, play an important role [25–27]. Variations in amino acid composition also reflect the ecological and functional adaptations of different taxa. For example, polyglutamine and polyalanine tracts are enriched in plant transcription factors, likely enhancing regulatory capacity [8]. Viral genomes, by contrast, are often optimized for minimal size and rapid replication, illustrating distinct constraints shaping protein composition [28].

Codon evolution may have been facilitated by broad specificity, meaning that a given codon may have encoded two or more amino acids. This phenomenon can still be observed in the case of the triplet UGA, which is a stop codon and encodes selenocysteine depending on the sequence context, as well as for other codons in some extant alternative codes [29]. For example, in the the *ascidian mitochondrial* code, the codons AGA and AGG can code for glycine, arginine or serine depending on specific circumstances [26,30,31].

At a certain point, however, the genetic code became “frozen” [10,21]. An important question is whether the result can be considered as a frozen accident, as suggested by Francis Crick [11,21] or was affected by the above-mentioned constraints [16]. According to the view of chance and necessity, both random and selective factors were relevant [32].

In a second phase of evolution, as life diversified and environmental conditions shifted, selective pressures on genomes changed as well, affecting the demand for particular amino acids and the costs of their synthesis. One of the most striking examples is methionine with a high biosynthetic cost and encoded by only one triplet in 22 of the 27 known genetic codes, acting as the initiator amino acid of proteins in almost all life forms and additionally occurring within proteins at non-start positions [4,33–35]. This amino acid shows a higher abundance in protein-coding genes than its encoding by one triplet would imply [7,22]. This may be related to the excision of the initial methionine in many proteins by post-translational modification [36]. Reconfiguring the codon-amino acid assignments of the genetic code in response to such pressures was likely to be unfeasible, given the fundamental constraints imposed by the frozen code [10,21]. Therefore, genomes evolved to modulate the relative abundances of amino acids in proteomes, thereby partially deviating from the initial correlation [19,37]. If these deviations are considerably high, two scenarios are conceivable, which are difficult to distinguish: Either the above-mentioned working hypothesis holds true for the first phase and the correlation between codon degeneracy and amino acid frequencies has disappeared in the second phase, or the hypothesis does not hold.

To some extent, the question under study is a chicken-and-egg problem: It is not easy to distinguish the cause from the effect. Has codon degeneracy evolved as a result of the required amino acid abundance or is the latter a result of the different number of synonymous codons? It has been hypothesized that the genetic code’s structure is optimized to minimize translational errors, with codons for chemically similar amino acids being separated by a single-nucleotide substitution (especially at the third position) thereby reducing the phenotypic impact of misreading or mutations [13,14,21,38]. These features align with error minimization theory, where the structure of the code enhances translation fidelity. Moreover, several of the genetic codes exhibit striking symmetry properties [3].

In this study, we empirically determine amino acid distributions in archaea, bacteria, eukaryotes, and viruses from protein-coding genes without taking into account differences in gene expression and post-translational modification such as the excision of the initial methionine in some proteins. To that end, sequence data of more than 20,000 proteomes available from the UniProt Knowledgebase [39] are analyzed. Thereafter, we examine the degree of correlation between these distributions and the theoretical predictions based on the numbers of synonymous codons in the standard genetic code as well as in several alternative genetic codes. This is be done with and without taking into account GC contents [26]. Additionally, we identify which amino acids adhere to the expected codon-based distributions most closely and which deviate.

## Materials and Methods

### Sequence and Genetic Code Data

We retrieve the nucleotide sequences of protein-coding genes and their corresponding amino acid sequences in FASTA format for all available archaea, bacteria, eukaryotes, and viruses from the UniProt Knowledgebase ([39], accessed on 13 July 2026, Table 1) using a custom Python script (v3.14). For simplicity, we consider all viruses in the data set as one additional domain. Note that some of the protein sequences in UniProt had been automatically translated from the genome sequence while others had been experimentally verified in proteomes. The UniProt and Taxonomy IDs for all organisms and sequences analyzed are listed in the Supplementary. The associated genetic codes for each organism are taken from the NCBI taxonomy browser [40,41] (Table 2). The bacterial, archaeal and plant plastid code (No. 11 in Table 1) only differs in the start codons (for a minority of proteins), while the encoded amino acids are the same as in the standard code. Therefore, for our analysis, codes 1 and 11 do not differ. Mapping tables for the genetic codes are obtained from the NCBI genetic code database [35]. If no associated genetic code is found for a species, the standard code (1) is used.

**Table 1.**
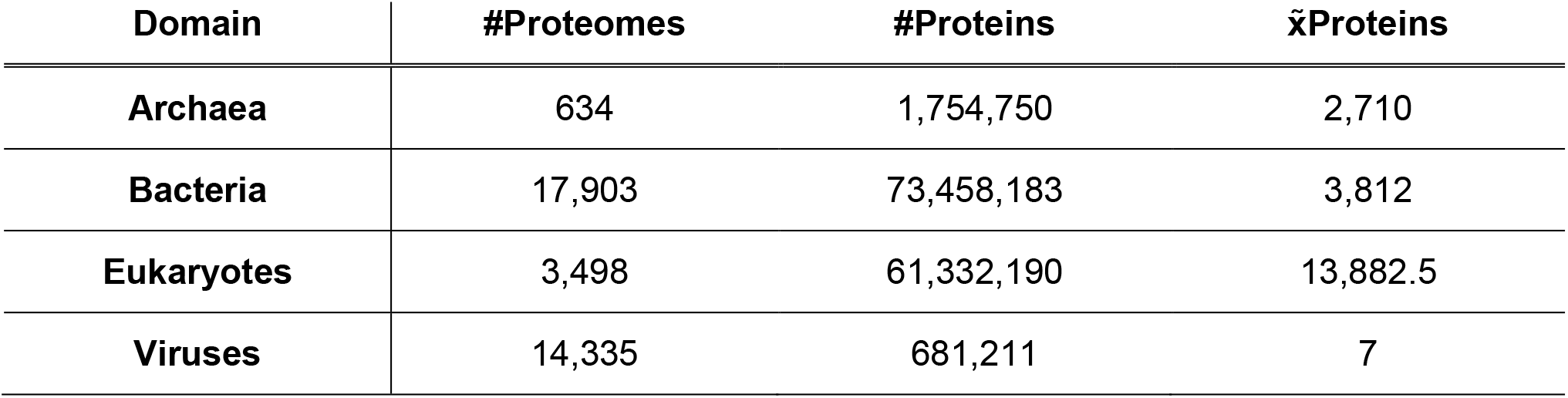
Distribution of proteomes and proteins (count and median) for each domain as available from the UniProt Knowledgebase [39].

**Table 2.**
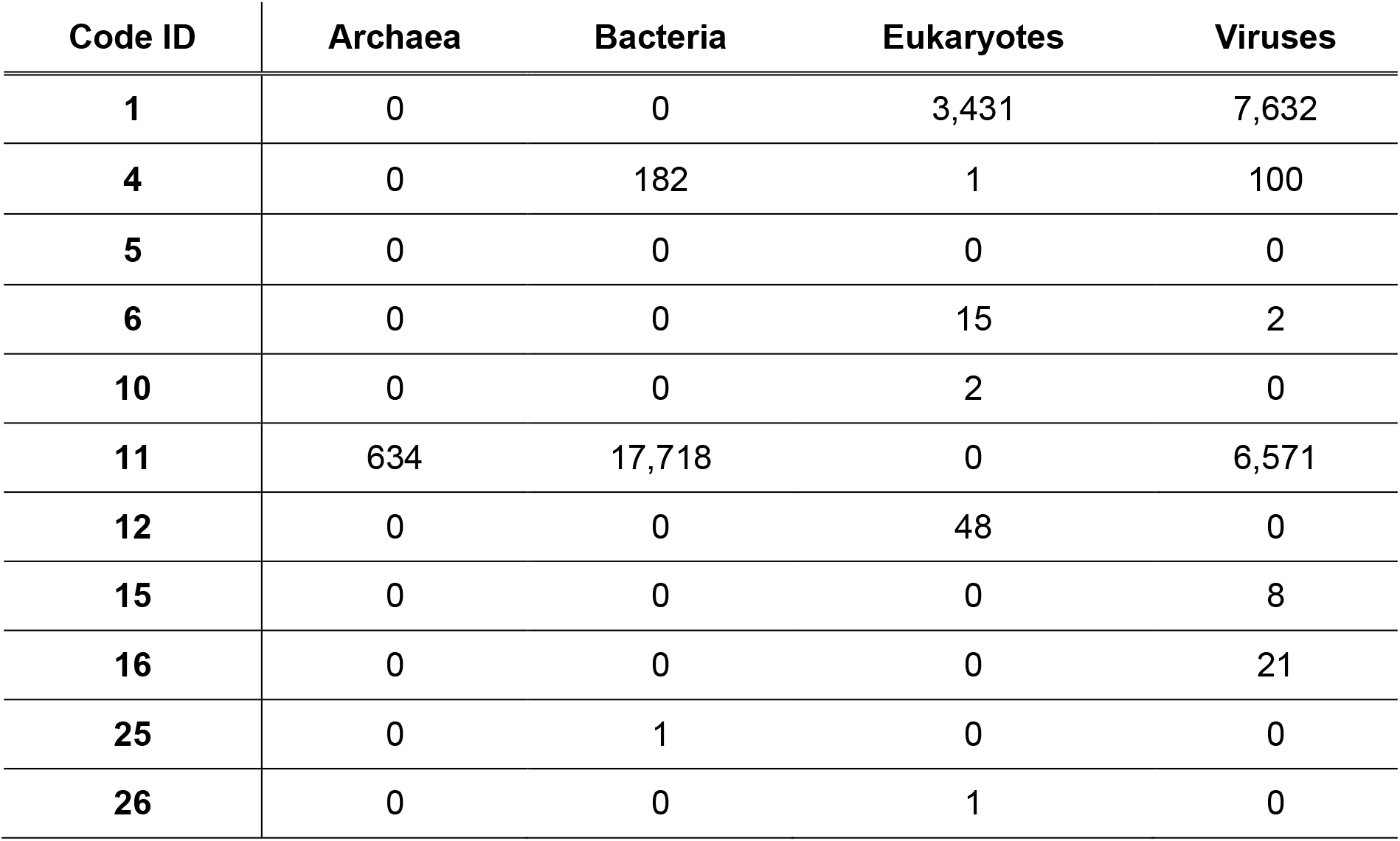
Distribution of genetic codes extracted for each domain from the NCBI taxonomy browser [40,41], with IDs sourced from the NCBI genetic code database [35].

### Empirical Amino Acid Distribution

We determine the relative amino acid frequency for the total set of protein-coding genes for each species across all domains. Differences in gene expression among proteins are not taken into account. Throughout this study, we refer to this unweighted protein set as the “genome-derived proteome”. Protein sequences are analyzed only if their corresponding gene sequences are also available, ensuring the inclusion of GC content for each protein. For each genome-derived proteome, amino acid frequencies were averaged by the median across all proteins and normalized so they sum to one.

### Theoretical Amino Acid Distribution

We determine the theoretical frequency of each amino acid by dividing its number of synonymous codons (in the genetic code under consideration) by the total number of sense codons. To account for the influence of GC content of protein-coding genes (which may differ from the overall genomic GC content), we calculate the theoretical amino acid frequency based on both the genetic code and median GC content (of each genome-derived proteome), following our previously described approach [26]. In the following, a short overview of the used formulas is given.

Considering Chargaff’s second parity rule [42], we can predict the frequency of nucleotides as a function of the GC content *g* [25]:

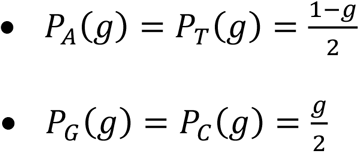

This parity rule does not hold in mitochondria, plastids, and several groups of viruses [43,44]. For simplicity, we disregard these exceptions. However, the formulas could be extended to three parameters, allowing all individual nucleotide frequencies to be accounted for by making one frequency dependent on the other three.

The predicted frequency of a triplet is then the product of all three nucleotides on the basis that each position is independent. For example, for the triplet CAT, we obtain:

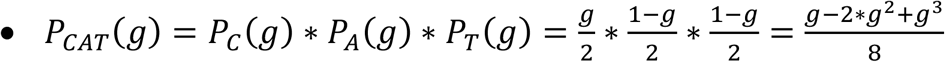

The predicted frequency of an amino acid is simply the sum of all synonymous triplets based on a given genetic code. For example, the equation for histidine in the standard genetic code reads:

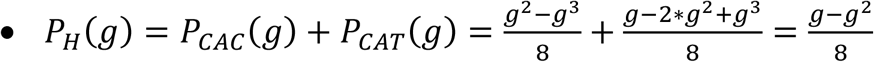

These calculations are here performed for all canonical amino acids and chosen genetic codes. In several alternative codes, for example, in the *invertebrate mitochondrial* code, the assignments of some triplets to amino acids are non-unique [45]. In that case, the triplet frequency is here evenly split between the respective amino acids, as done earlier [14].

### Correlation Calculations of Compositional Data

We compare the empirical amino acid frequencies of each genome-derived proteome within a domain to the theoretical frequencies derived from the associated genetic code, both with and without adjustments for GC content. However, these data sets are compositional [46]: the frequencies of the 20 amino acids within each proteome sum to one. Under this constraint, analyzing raw frequencies can induce spurious trade-offs (if one amino acid frequency increases, at least one other must decrease), potentially leading to artificial positive or negative correlations [47]. To avoid these artifacts, we work in log-ratio coordinates using the centered log-ratio (CLR) transform, which maps compositions to Euclidean space where distances and correlations are interpretable [48,49]. All comparisons are therefore performed on CLR-transformed values:

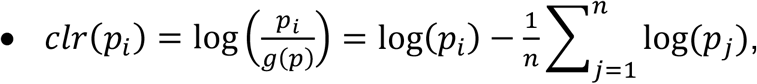

where *clr*(*p_i_*) is the centered log-ratio transformed value of amino acid *i*, *p_i_* the frequency of amino acid *i* and *g*(*p*) the geometric mean of the given frequencies. To prevent undefined values in logarithmic calculations, amino acid frequencies of zero were substituted with *e*^−12^. Deviations between observed amino acid frequencies of each genome-derived proteome and expected values (based on synonymous number of codons alone and adjusted for GC content) are quantified using the Aitchison distance [50]:

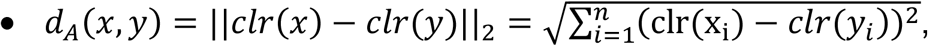

where *x* and *y* are two sets of frequencies.

For the correlation analysis, we applied Pearson (*r*), Spearman’s rank (ρ), and Kendall’s Tau (*τ*) correlation coefficient tests to the CLR-transformed values [51–53]. To ensure robustness of all tests, we conducted permutations tests with 100,000 iterations. Correlation coefficients and *p*-values were combined using Fisher’s Z-transformation and Fisher’s method, respectively [54,55].

### Balances of Codon Degeneracy Distributions

Additionally, we analyze the compositional data by an alternative method, which is to construct an orthonormal log-ratio basis based on sequential binary partitions (SBP) [49,56]. An SBP calculates a so-called balance or contrast between two distinct subgroups of data given the following equation, in which the argument of the logarithm is the quotient of two geometric means [57]:

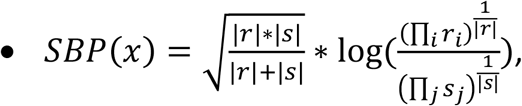

where *r* and *s* are two distinct groups of a frequency set *x* based on a specific property. A positive balance assigns, on average, the group *r* a higher importance in the dataset than the group *s*, and vice versa for a negative balance. A log-ratio of zero implies that the two sets of amino acids occur equally often. Subsequently, *r* and *s* are further recursively split into two distinct groups and their balances are calculated until only singleton sets remain, leading to *D* − 1 balances (where *D* is the size of the input set). In our case, we are interested in a subset of properties, namely the balances between the frequencies of high-codon amino acids and low-codon amino acids. It is worth noting that a recursive decomposition of the set of amino acids has been applied earlier [58].

Using the standard genetic code, we divide amino acids (and their frequencies) into sets with at least three codons and amino acids with at most two codons. The first group is further divided into amino acids with four and six codons and amino acids with three codons. This is repeated until only frequency sets with a single number of codons remain, resulting in four distinct balances and five single property sets (Table 3).

**Table 3.**
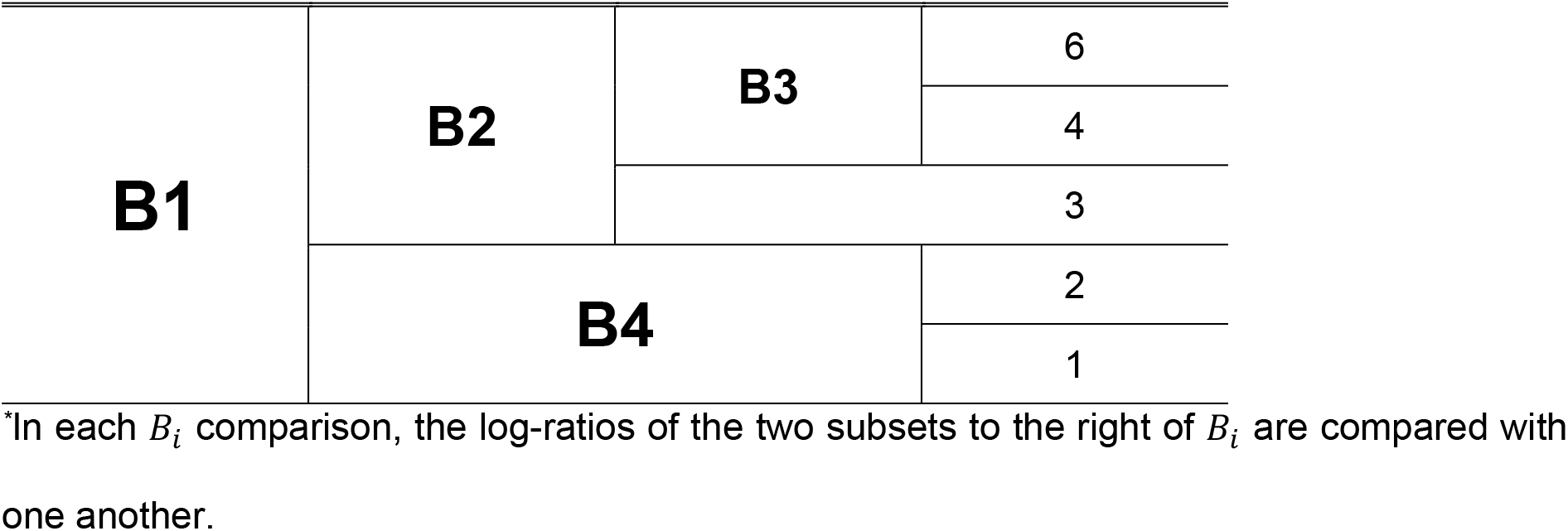
The recursive splits for the calculations of balances (B1 to B4) based on the different number of codons in the standard genetic code*.

### Integer Linear Programming of Codon Degeneracy

Lastly, the problem described here can also be viewed in reverse. Instead of examining how close the observed frequencies of each amino acid are to their codon distributions, one can investigate what the optimal number of codons for each amino acid would be given their observed frequencies. We have to consider that the number of synonymous codons represent compositional and integer (i.e. natural number) data. In our case, this leads to an integer linear programming (ILP) problem [59]. A plausible objective function is to minimize the following sum of differences:

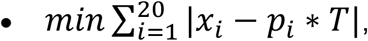

where *x_i_* is the optimal number of synonymous codons for amino acid *i*, *p_i_* is the observed frequency of amino acid *i*, and *T* is the available number of overall codons. *x_i_* and *T* are further constraint as follows:

• *x_i_* ≥ 1, *x* ∈ ℤ, ∀*i*

• *T* = 64 − #*Stops_g_*

Here, each allocation of synonymous number of codons *x_i_* must be a positive integer and at least one (so that each amino acid receives at least one codon). *T* corresponds to the number of all possible codons (i.e. 64) minus the number of stop codons associated with the genetic code *g* of the observed organism (so that the entire codon space is used). For example, in the standard genetic code, three codons are assigned to the stop signal, resulting in *T* = 61 available codons.

Overall deviations between the assigned codon degeneracy based on the genetic codes and the optimal codon degeneracy based on the observed frequencies are quantified using the Jensen-Shannon divergence [60]:

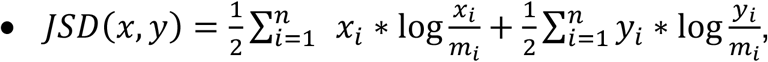

where *x* and *y* are two sets of codon degeneracy distributions and *m* is the midpoint mixture distribution between the two sets:

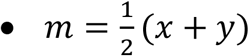

This divergence measure has the advantage that it is symmetric, unlike the Kullback-Leibler divergence [61].

## Results

### Comparative Analysis of GC Content across Domains

The GC content corresponding to the genome-derived proteome exhibits distinct patterns across different domains, with bacteria and archaea showing higher GC contents and greater variations than eukaryotes and viruses. Bacteria exhibit the highest median of 57.64 % (±11.66 %; Figure 1), closely followed by archaea with a median of 56.79 % (±10.69 %). In comparison, eukaryotes and viruses exhibit lower GC content medians and variability of 50.98 % (±4.07 %) and 46.15 % (±5.69 %) respectively.

**Figure 1.**
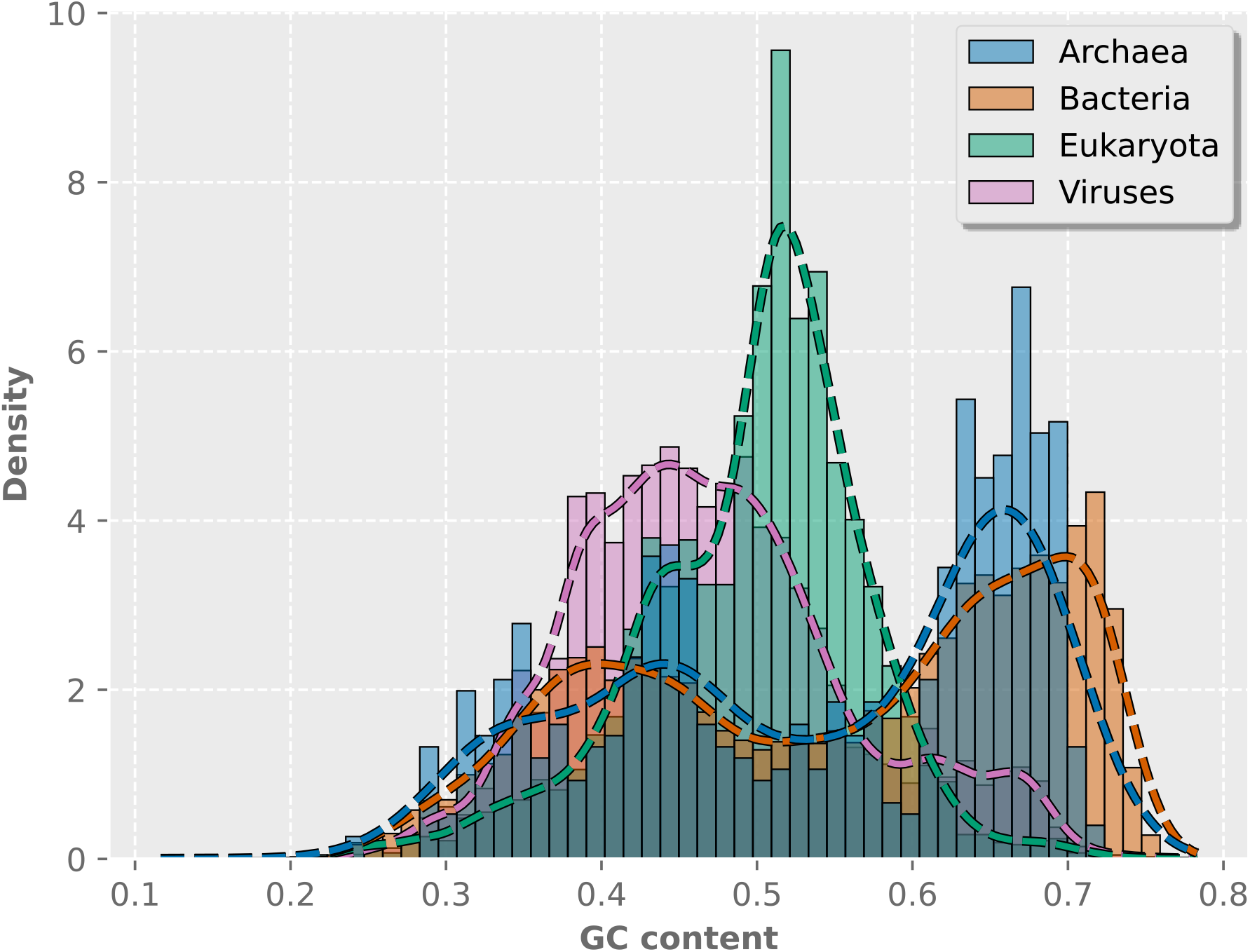
Density distribution of median GC content of protein-coding genes per species across the four domains. Blue, archaea; orange, bacteria; green, eukaryotes; pink, viruses. The dashed curves show kernel density estimates that smooth each histogram. The number of bins is set to ten.

The data on protein number and the distribution of protein lengths can be found in the Supplementary. These are not directly relevant to our study.

### Amino Acid Abundances and Variability across Domains

In the following paragraphs, we use the names of amino acids as a representative for the set of triplets encoding them. Overall, amino acid distributions are largely consistent across domains, with variations in median abundances and variability for specific amino acids (Figure 2, Table 4). Leucine (that is, the set of its six codons) is the most abundant amino acid in archaea, eukaryotes, and viruses. Alanine is the most abundant amino acid in bacteria. Tryptophan, cysteine and histidine are consistently the least frequent amino acids across all domains.

**Figure 2.**
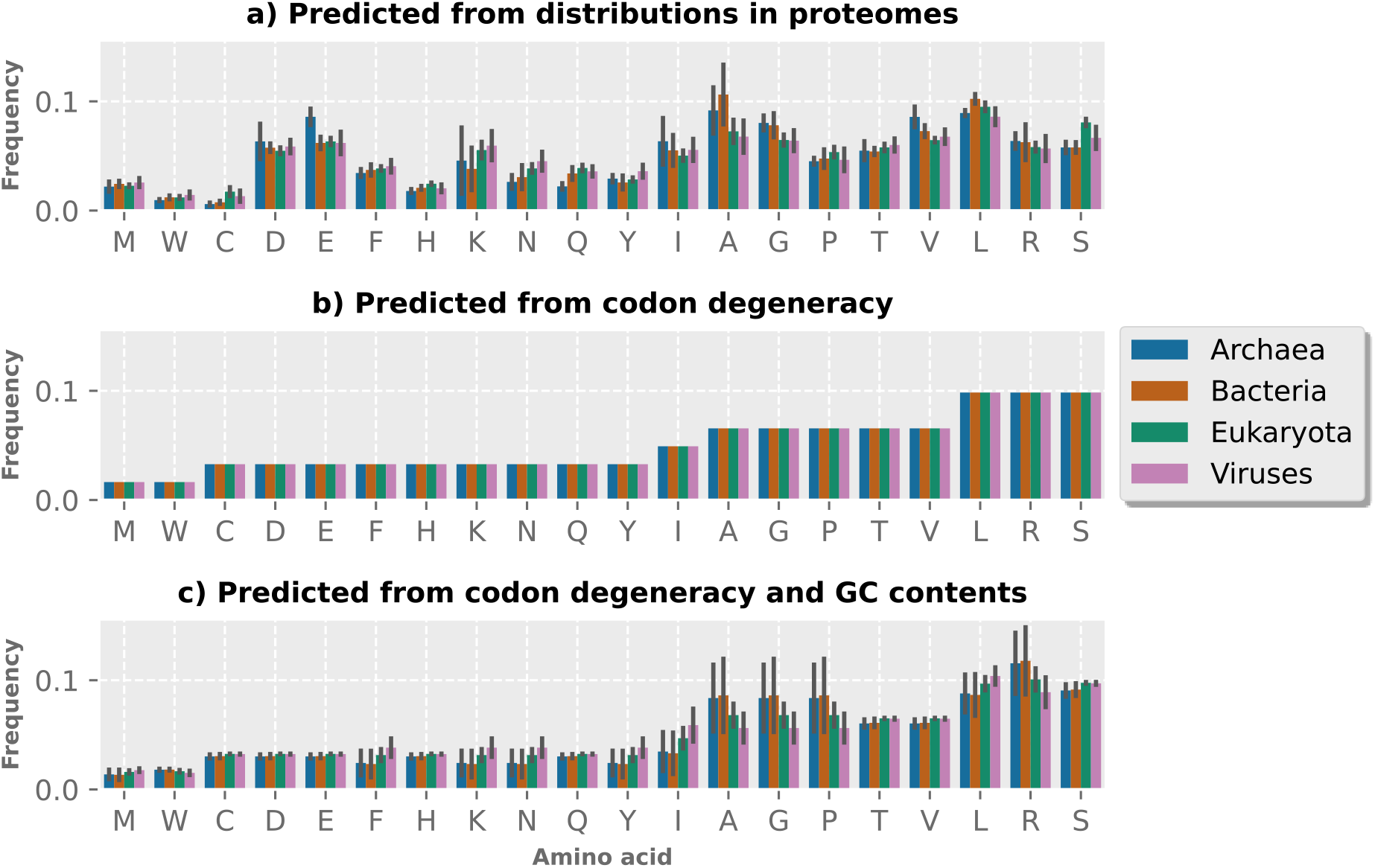
Median frequencies of amino acids and their median absolute deviations across the four domains. a) Determined empirically from the genomic data, b) predicted from codon degeneracy given the associated genetic code, and c) predicted from the codon degeneracy given the associated genetic code adjusted for GC contents. Blue, archaea; orange, bacteria; green, eukaryotes; pink, viruses.

**Table 4.**
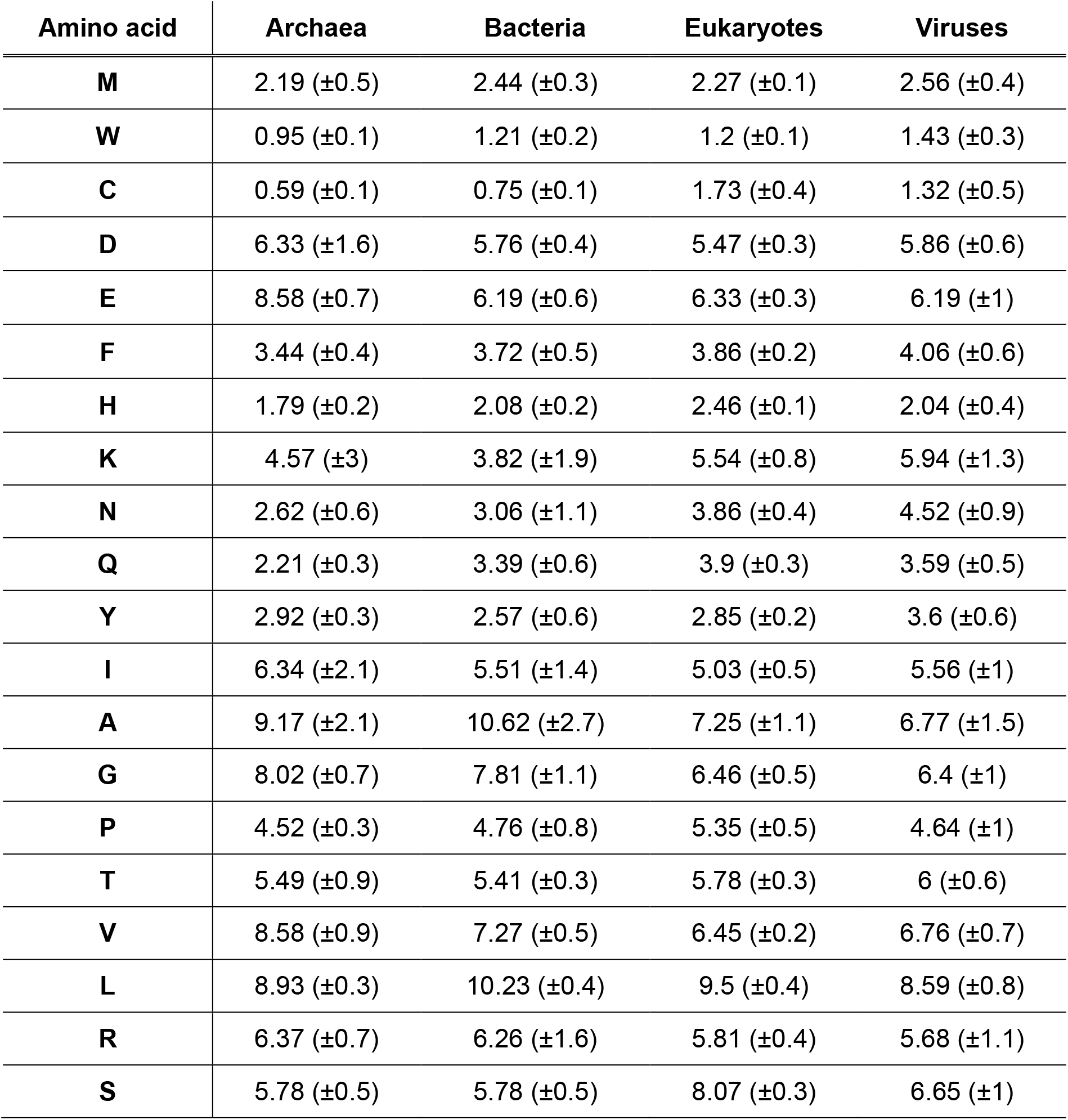
Observed median distributions of amino acids (ordered as in Figure 2) and their median absolute deviations (in %) in genome-derived proteome across domains.

| Amino acid | Archaea | Bacteria | Eukaryotes | Viruses |
| --- | --- | --- | --- | --- |
| **M** | 2.19 ($\pm 0.5$) | 2.44 ($\pm 0.3$) | 2.27 ($\pm 0.1$) | 2.56 ($\pm 0.4$) |
| **W** | 0.95 ($\pm 0.1$) | 1.21 ($\pm 0.2$) | 1.2 ($\pm 0.1$) | 1.43 ($\pm 0.3$) |
| **C** | 0.59 ($\pm 0.1$) | 0.75 ($\pm 0.1$) | 1.73 ($\pm 0.4$) | 1.32 ($\pm 0.5$) |
| **D** | 6.33 ($\pm 1.6$) | 5.76 ($\pm 0.4$) | 5.47 ($\pm 0.3$) | 5.86 ($\pm 0.6$) |
| **E** | 8.58 ($\pm 0.7$) | 6.19 ($\pm 0.6$) | 6.33 ($\pm 0.3$) | 6.19 ($\pm 1$) |
| **F** | 3.44 ($\pm 0.4$) | 3.72 ($\pm 0.5$) | 3.86 ($\pm 0.2$) | 4.06 ($\pm 0.6$) |
| **H** | 1.79 ($\pm 0.2$) | 2.08 ($\pm 0.2$) | 2.46 ($\pm 0.1$) | 2.04 ($\pm 0.4$) |
| **K** | 4.57 ($\pm 3$) | 3.82 ($\pm 1.9$) | 5.54 ($\pm 0.8$) | 5.94 ($\pm 1.3$) |
| **N** | 2.62 ($\pm 0.6$) | 3.06 ($\pm 1.1$) | 3.86 ($\pm 0.4$) | 4.52 ($\pm 0.9$) |
| **Q** | 2.21 ($\pm 0.3$) | 3.39 ($\pm 0.6$) | 3.9 ($\pm 0.3$) | 3.59 ($\pm 0.5$) |
| **Y** | 2.92 ($\pm 0.3$) | 2.57 ($\pm 0.6$) | 2.85 ($\pm 0.2$) | 3.6 ($\pm 0.6$) |
| **I** | 6.34 ($\pm 2.1$) | 5.51 ($\pm 1.4$) | 5.03 ($\pm 0.5$) | 5.56 ($\pm 1$) |
| **A** | 9.17 ($\pm 2.1$) | 10.62 ($\pm 2.7$) | 7.25 ($\pm 1.1$) | 6.77 ($\pm 1.5$) |
| **G** | 8.02 ($\pm 0.7$) | 7.81 ($\pm 1.1$) | 6.46 ($\pm 0.5$) | 6.4 ($\pm 1$) |
| **P** | 4.52 ($\pm 0.3$) | 4.76 ($\pm 0.8$) | 5.35 ($\pm 0.5$) | 4.64 ($\pm 1$) |
| **T** | 5.49 ($\pm 0.9$) | 5.41 ($\pm 0.3$) | 5.78 ($\pm 0.3$) | 6 ($\pm 0.6$) |
| **V** | 8.58 ($\pm 0.9$) | 7.27 ($\pm 0.5$) | 6.45 ($\pm 0.2$) | 6.76 ($\pm 0.7$) |
| **L** | 8.93 ($\pm 0.3$) | 10.23 ($\pm 0.4$) | 9.5 ($\pm 0.4$) | 8.59 ($\pm 0.8$) |
| **R** | 6.37 ($\pm 0.7$) | 6.26 ($\pm 1.6$) | 5.81 ($\pm 0.4$) | 5.68 ($\pm 1.1$) |
| **S** | 5.78 ($\pm 0.5$) | 5.78 ($\pm 0.5$) | 8.07 ($\pm 0.3$) | 6.65 ($\pm 1$) |

In the comparison among domains, aspartate and glutamate (used here synonymously with aspartic acid and glutamic acid, respectively) occur at elevated levels in archaea. Lysine, asparagine and glutamine are more abundant in eukaryotes and viruses. Alanine and glycine are found in higher proportions in archaea and bacteria. Tyrosine is slightly elevated in viruses. Proline occurs slightly more frequently in eukaryotes, while valine shows elevated levels in bacteria and, especially, archaea. Serine is more prevalent in eukaryotes and viruses. Despite these variations, the overall amino acid distributions exhibit broad similarities across biological domains, with leucine, alanine, and glycine among the most common.

### Correlation to Synthesis Costs

To resolve the chicken-and-egg problem mentioned in the Introduction, it is of interest to speculate about the initial cause for the frequency distribution of amino acids: Did amino acid frequencies influence the number of assigned synonymous codons or vice versa? It is a straightforward assumption that the distribution reflects a balance between the need for functional diversity and the metabolic costs of amino acid synthesis, where simpler and less costly amino acids dominate proteomes [4,33]. Costs in metabolism can be quantified by the moles of substrate (e.g. glucose) to produce one mole of amino acid. It is plausible that abundance and metabolic amino acid-over-substrate yield are correlated [5]. That means that abundance and cost are inversely correlated.

Table 5 and Figure 3 show the calculated molar yield for the special case of *Escherichia coli* (taxonomy ID: 83333) for three initial substrates (glucose, glycerol and acetate) compared to observed amino acid distributions as well as their correlation coefficients [33]. As can be seen, moderate correlations exist for all substrates (glucose: *r* = 0.43, ρ = 0.44, *τ* = 0.3; glycerol: *r* = 0.39, ρ = 0.4, *τ* = 0.27; acetate: *r* = 0.45, ρ = 0.48, *τ* = 0.33). The tests were, however, only significant (*p* < 0.05) for acetate as the initial substrate.

**Table 5.**
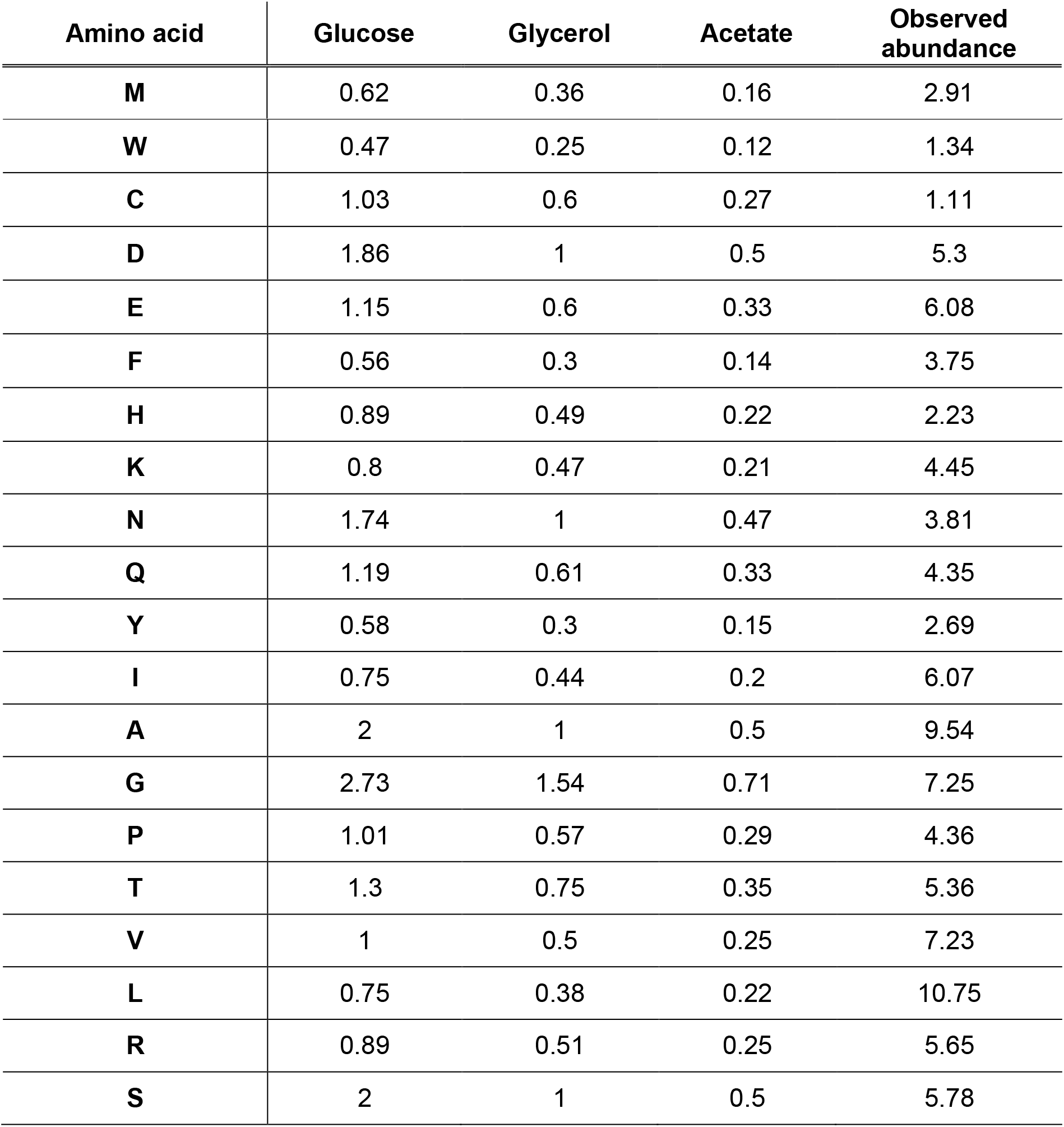
Biosynthesis yields (mol/mol) for each amino acid (ordered as in Figure 2) in E. *coli* (taxonomy ID: 83333) from glucose, glycerol, and acetate computed by the linear programming standard approach [33] as well as median amino acid distribution as determined in our analysis (in %).

**Figure 3.**
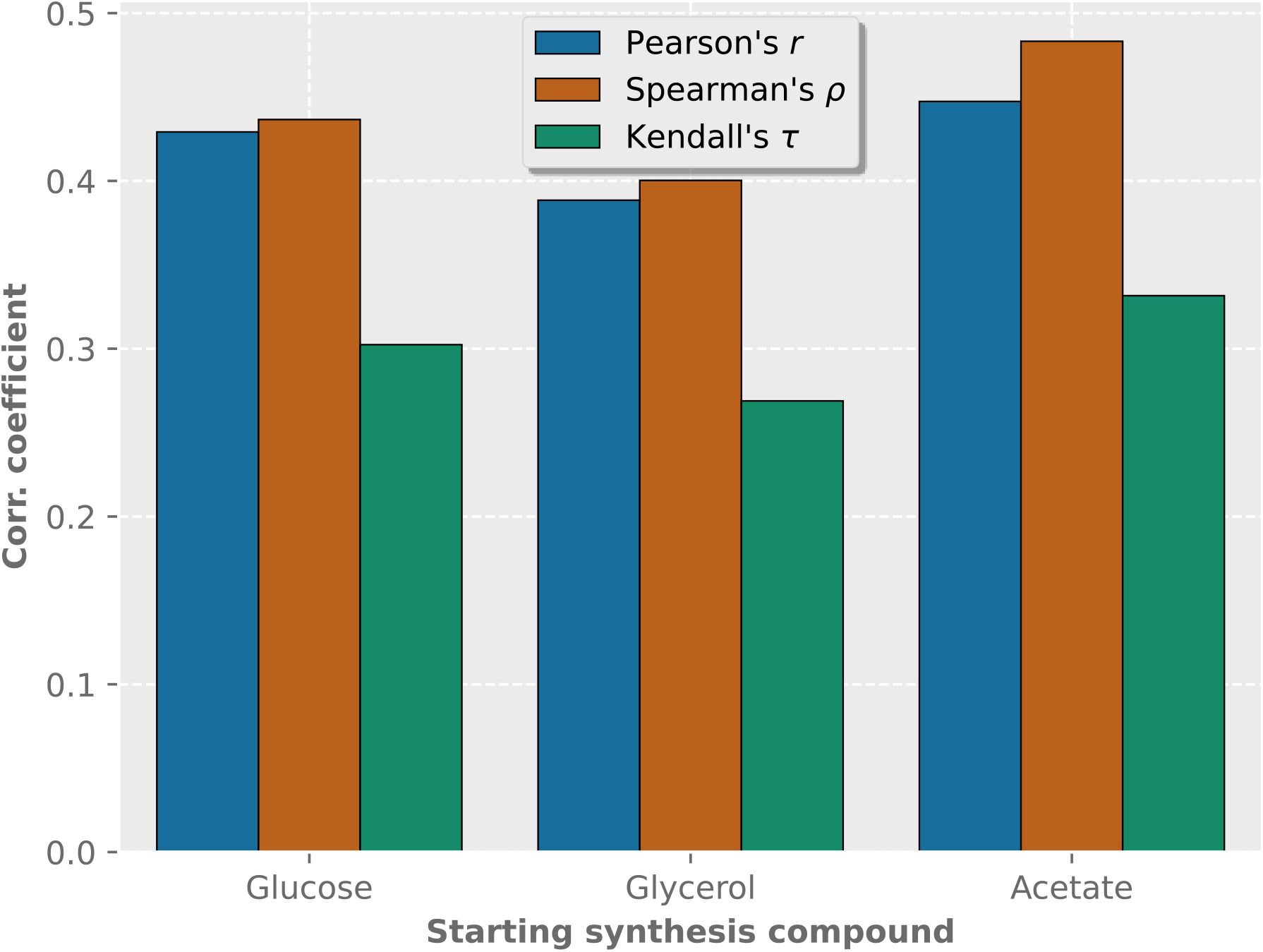
Correlation coefficients between median amino acid distributions and biosynthesis yields (mol/mol, linear programming standard approach) for each amino acid in *E. coli* (taxonomy ID: 83333) from glucose, glycerol, and acetate [33]. Blue, Pearson’s *r*; orange, Spearman’s *ρ*; green, Kendall’s *τ*.

It is important to note that molar amino acid yields vary considerably across species. In particular, several amino acids are essential in humans and many other animals, as they cannot be synthesized internally. As a result, quantifying their individual yields or costs is challenging. Given these complexities, we do not elaborate on metabolic yields or costs in this study, focusing instead on the correlation between amino acid frequency and codon multiplicity.

### Deviations between Observed and Theoretical Amino Acid Distributions across Domains

Next, the median amino acid distributions of archaea, bacteria, eukaryotes, and viruses are compared to theoretical distributions based on codon degeneracy alone and codon degeneracy adjusted for GC content (Figure 4, Table 6). The comparisons are made in centered log-ratio (CLR) space to avoid spurious effects. Based on codon degeneracy, methionine, glutamate, aspartate and lysine (in non-bacterial proteomes) consistently show positive deviations as well as alanine in the domains of archaea and bacteria. In contrast, negative deviations are observed for cysteine, histidine, proline, tryptophan and arginine as well as serine in non-eukaryotic domains and glutamine in archaeal proteomes. Adjustment for GC content generally yields similar patterns, with a few exceptions. Isoleucine and phenylalanine show higher positive deviations in archaea and bacteria, while alanine tends to match more closely the observed frequency.

**Figure 4.**
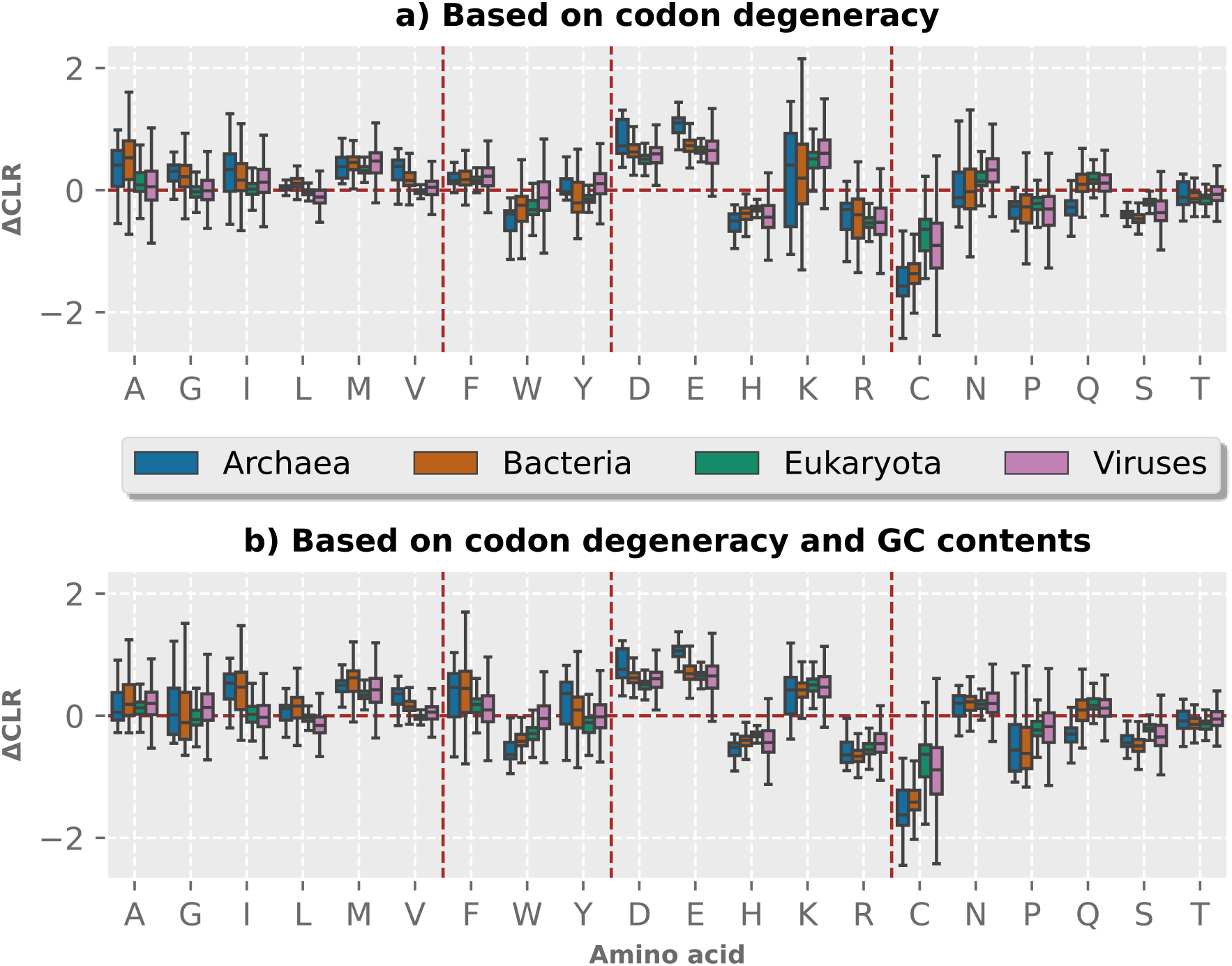
Individual deviations between empirical values and theoretical values for the 20 canonical amino acids in CLR space. a) Derived from codon degeneracy, b) derived from codon degeneracy adjusted for GC content. Outliers are excluded to improve visualization. The horizontal brown lines indicate zero deviations. Vertical brown lines separate amino acids into the following categories (in that order): Aliphatic, aromatic, charged, and uncharged. Blue, archaea; orange, bacteria; green, eukaryotes; pink, viruses.

**Table 6.**
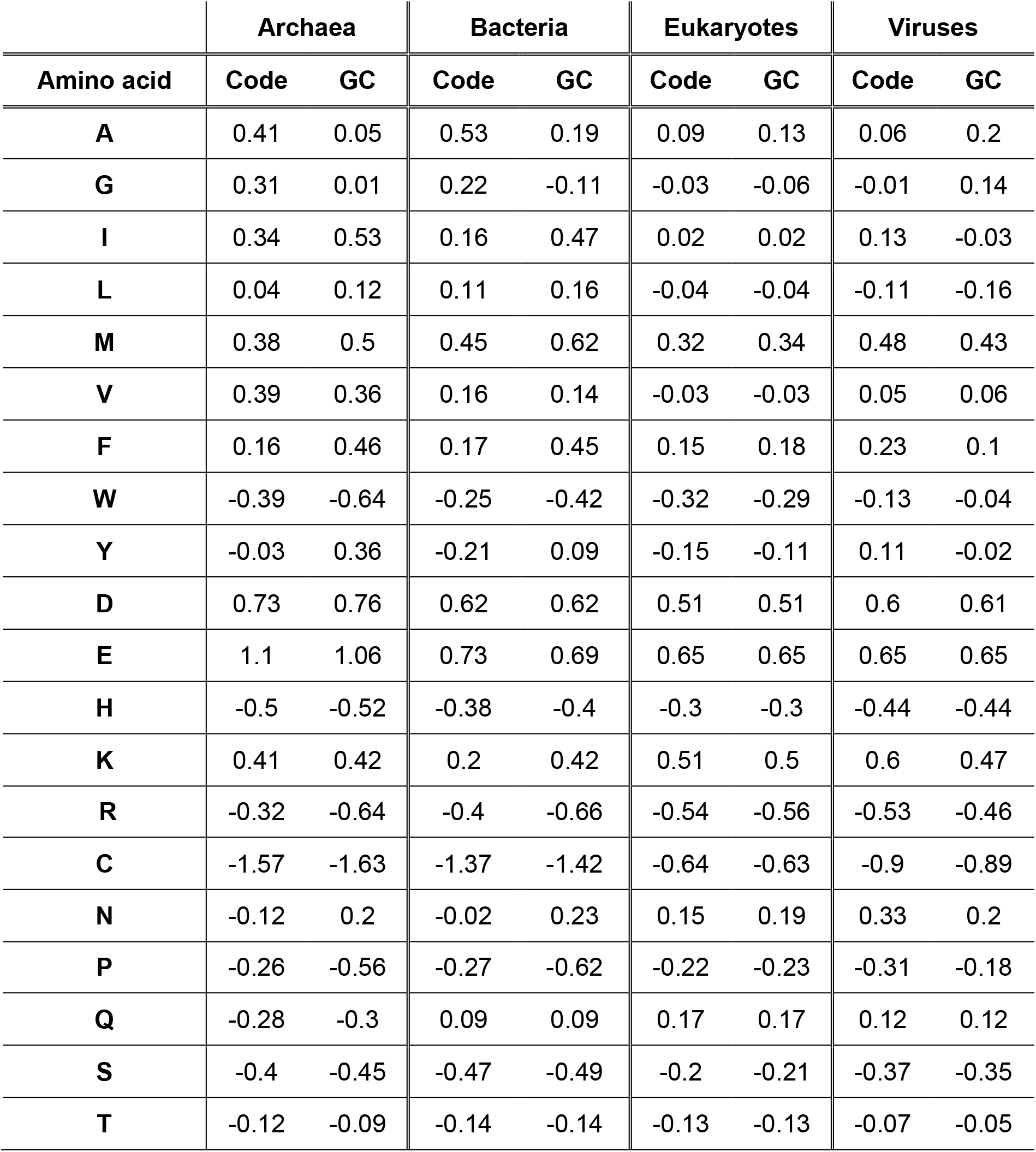
Median deviations between empirical values and theoretical values derived from codon degeneracy and from codon degeneracy adjusted for GC content for the 20 canonical amino acids in CLR space (ordered as inFigure 4).

Looking at the overall median Aitchison distances between observed and expected amino acid distributions, the values generally decrease from archaea (Codon: 2.71, GC: 2.84) over bacteria (Codon: 2.31, GC: 2.5) and viruses (Codon: 2.07, GC: 1.96) to eukaryotes (Codon: 1.58, GC: 1.6) (Figure 5).

**Figure 5.**
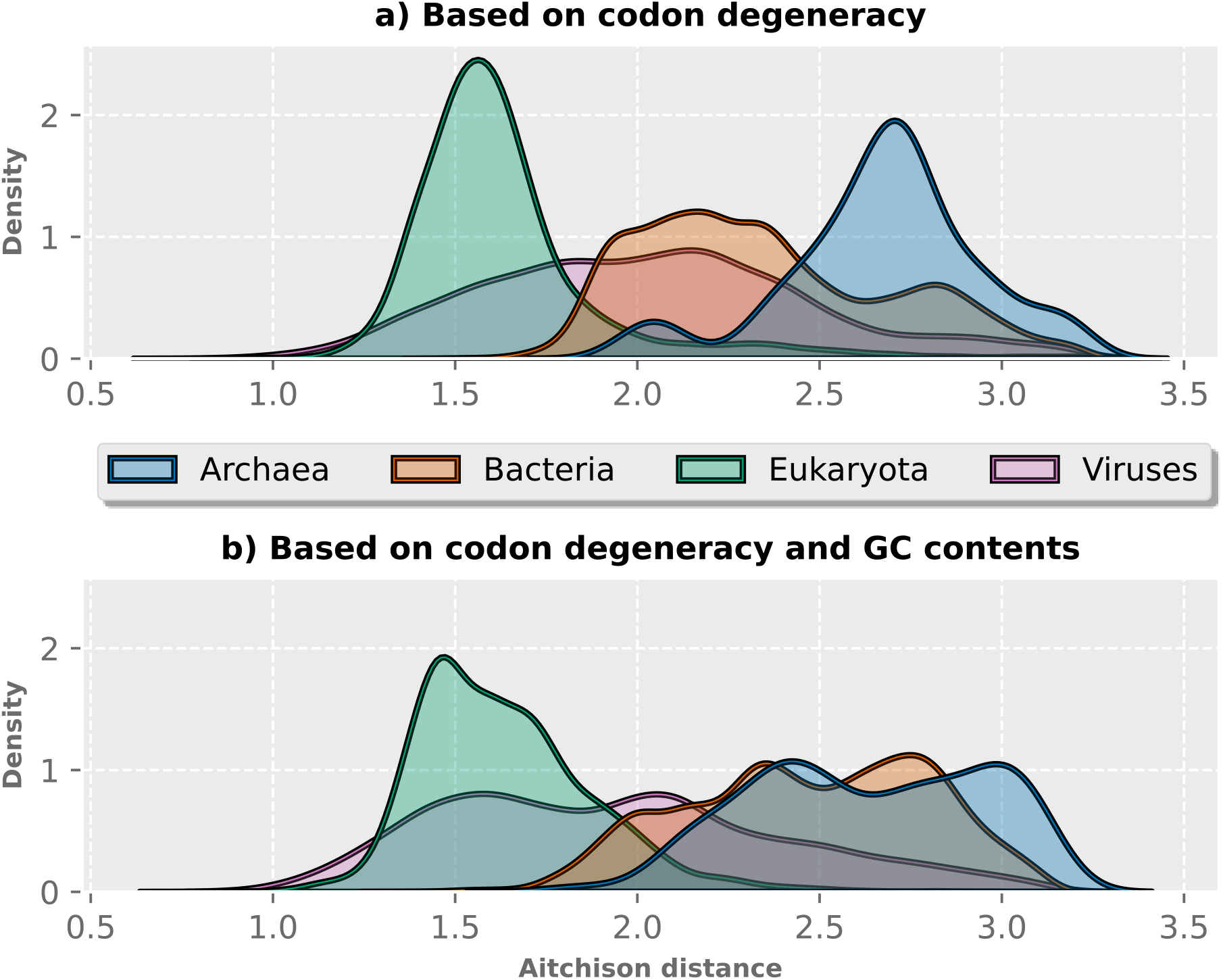
Aitchison distances between empirical values and theoretical values for the 20 canonical amino acids in CLR space. a) Derived from codon degeneracy, b) derived from codon degeneracy adjusted for GC content. Distances above the 95^th^ percentile are excluded to improve visualization. Blue, archaea; orange, bacteria; green, eukaryotes; pink, viruses.

### Correlations: Trends in Codon Usage and GC Content Influence

The correlations between empirical amino acid distributions and theoretical expectations based on codon multiplicity and GC content adjustment were assessed across the four domains using Pearson, Spearman’s rank and Kendall’s Tau correlation coefficient tests (Figure 6). The tests are done in CLR space to avoid spurious effects. The resulting average coefficients follow a consistent pattern, both with and without adjustments for GC content, with archaea exhibiting the lowest correlations (*r* = 0.64, ρ = 0.62, *τ* = 0.49), followed by bacteria (*r* = 0.67, ρ = 0.69, *τ* = 0.55) and viruses (*r* = 0.66, ρ = 0.68, *τ* = 0.54), while eukaryotes (*r* = 0.77, ρ = 0.78, *τ* = 0.66) show the highest correlations. Overall, adjusting for GC content shows narrower density distributions for all correlations across all domains under study, with elevated minimum values while reducing maximum values, resulting in a higher and sharper central peak (see Supplementary). Consequently, this results in only minor changes in overall correlation coefficients, indicating that the number of synonymous codons for each amino acid alone already provides a strong predictive model for amino acid distributions across genome-derived proteomes. All correlation tests returned a *p*-value below 0.05 on average (see Supplementary).

**Figure 6.**
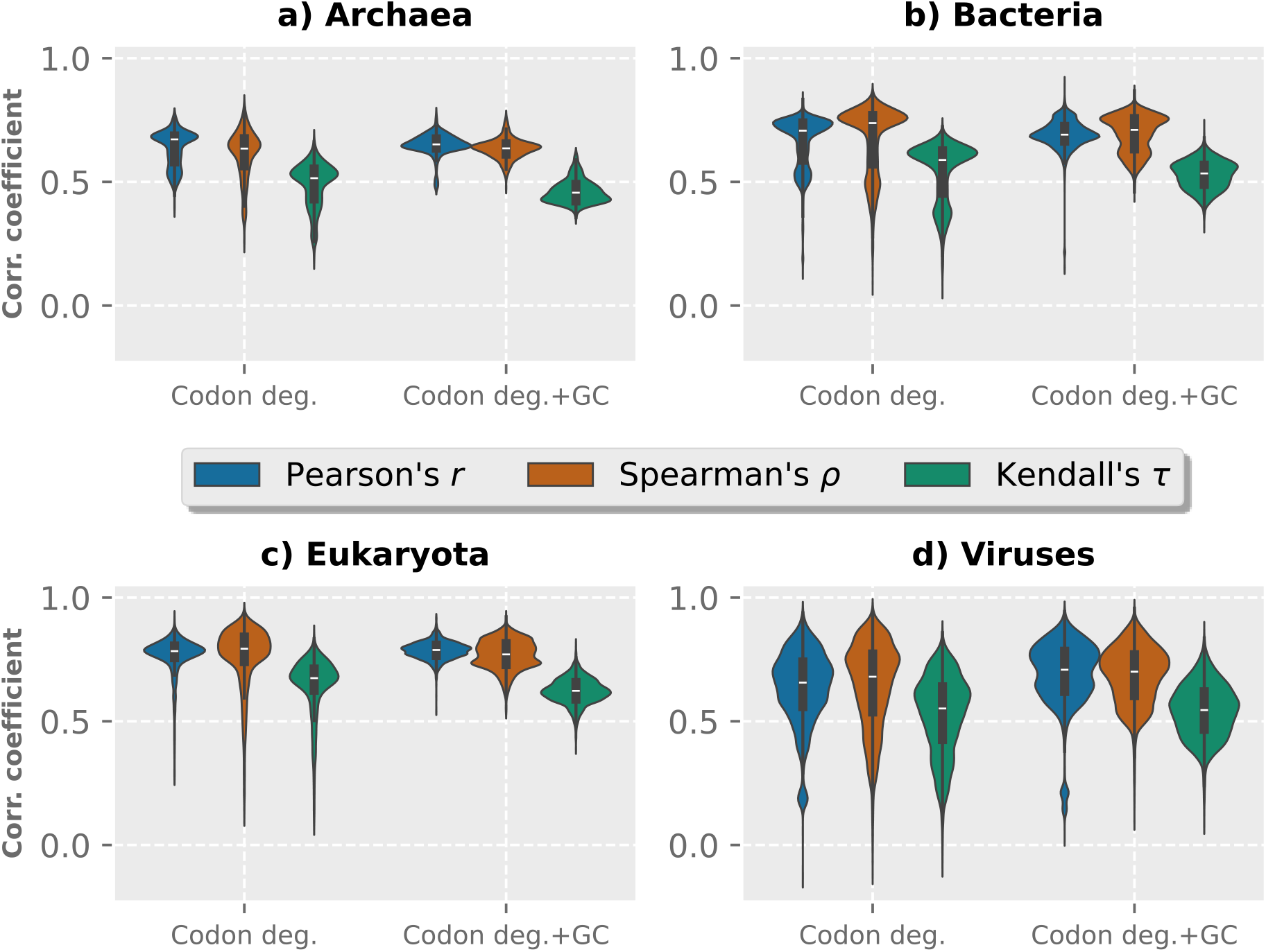
Correlation coefficients between empirical amino acid distributions per species and theoretical expectations based on codon degeneracy and codon degeneracy adjusted for GC content across the four domains. Blue, Pearson’s *r*; orange, Spearman’s *ρ*; green, Kendall’s *τ*.

### Distributions of Positively and Negatively Charged Amino Acid

Next, we examine the distributions of positively charged amino acids (histidine, lysine, arginine) and negatively charged ones (glutamate, aspartate) at near-neutral pH (Figure 7). It is striking that all five charged amino acids show notable deviations in their abundance from the predictions based on codon degeneracy. It is worth noting that at any pH value, the total charge of all amino acids need not be zero. Proteins can have a total charge, which causes, for example, a Donnan potential across membranes [62,63]. Interestingly, the overall frequencies of both groups remain relatively stable across a wide range of GC contents. On average, positively charged amino acids occur more frequently than negatively charged ones in eukaryotes (positive: 13.8 %, negative: 11.81 %) and viruses (positive: 13.66 %, negative: 12.05 %). In archaea, by contrast, negatively charged amino acids are more abundant (positive: 12.73 %, negative: 14.92 %). In bacteria, the two groups occur at nearly equal frequencies (positive: 12.16 %, negative: 11.95 %).

**Figure 7.**
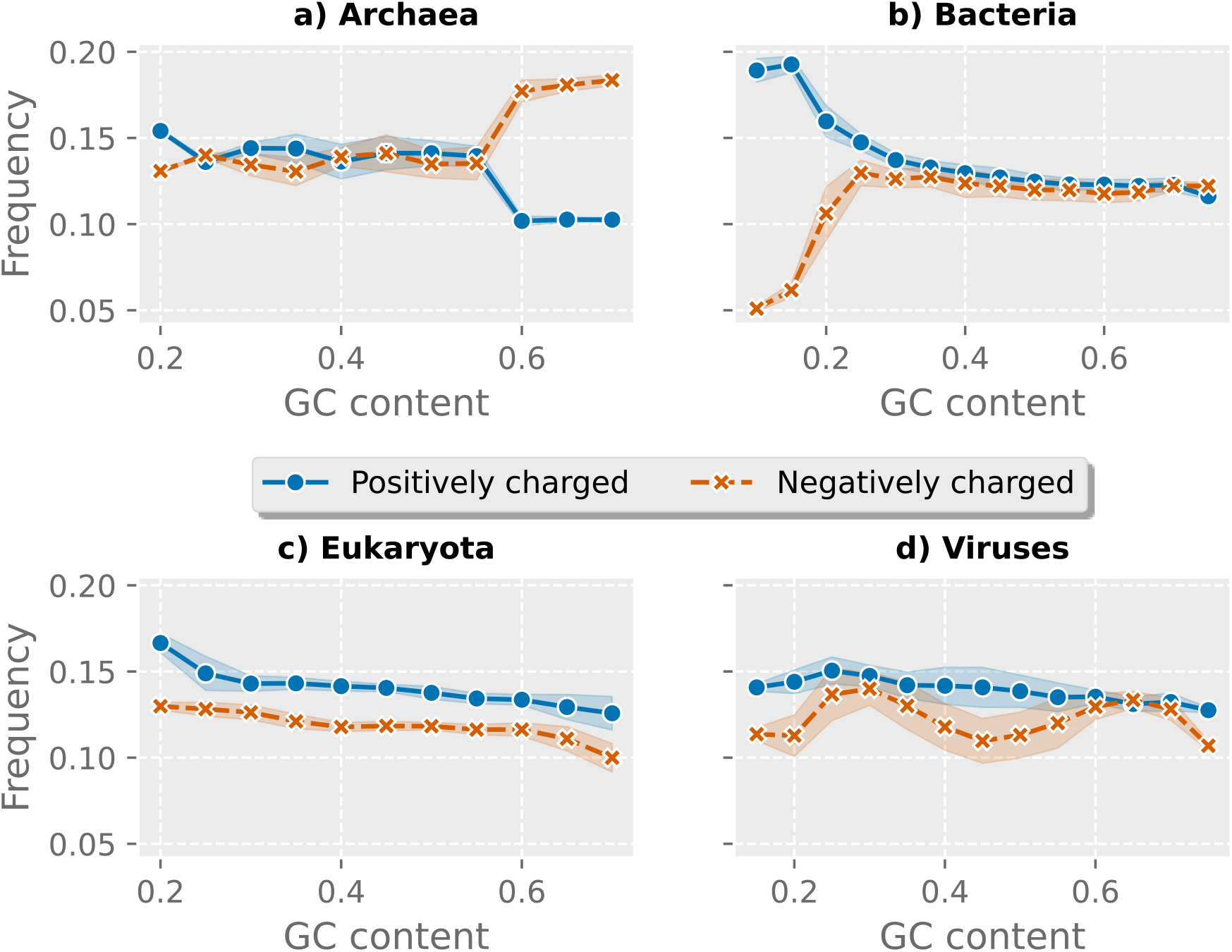
Median distributions of positively (blue) and negatively (orange) charged amino acids (histidine, lysine, arginine vs. glutamate, aspartate) across the four domains with median absolute deviations indicated, with GC content on the *x*-axis in 5 % intervals.

While there is generally no significant correlation between the dependencies of these two groups of amino acids on GC content, bacteria with low GC content tend to have more positively charged and fewer negatively charged amino acids (Figure 7a), whereas archaea with high GC content show the opposite pattern, with more negatively charged and fewer positively charged amino acids (Figure 7b).

### Balances: Preference of High-Codon Amino Acids

Next, we computed balance values based on the number of codons for each amino acid, as explained in the Methods section. Depending on the genetic code, different codon degeneracy sets and thus different balances are obtained. Here, we focus on the sets with codon multiplicity of one, two, three, four, and six (as in the standard genetic code), resulting in four balances for ease of interpretation. This distribution of synonymous number of codons is used by almost all organisms in the dataset, regardless of the domain (archaea: 100 %, bacteria, 98.98 %, eukaryotes: 98.08 %, viruses: 99.09 %).

Overall, the balance **B1** (i.e., the log-ratio between amino acids with at least three codons and amino acids with at most two codons) shows, on average, the highest abundance log-ratio (approximately 1.79) for each domain, indicating that amino acids with a high number of codons are preferred over amino acid with a low number of codons for incorporation into genome-derived proteomes (Figure 8). The second highest is balance **B4** (i.e., the log-ratio between two-codon amino acids and one-codon amino acids) with an average balance value half as high as **B1** (approximately 0.87). This is probably due, on the one hand, to the small number of amino acids with one codon (in the standard genetic code, only methionine and tryptophan) and, on the other hand, to the overabundance of glutamate and aspartate. Finally, balances **B2** and **B3** show values close to zero in all four domains (**B2**: 0.18, **B3**: 0.14), indicating that there is a minor difference between amino acids with three, four, and six codons.

**Figure 8.**
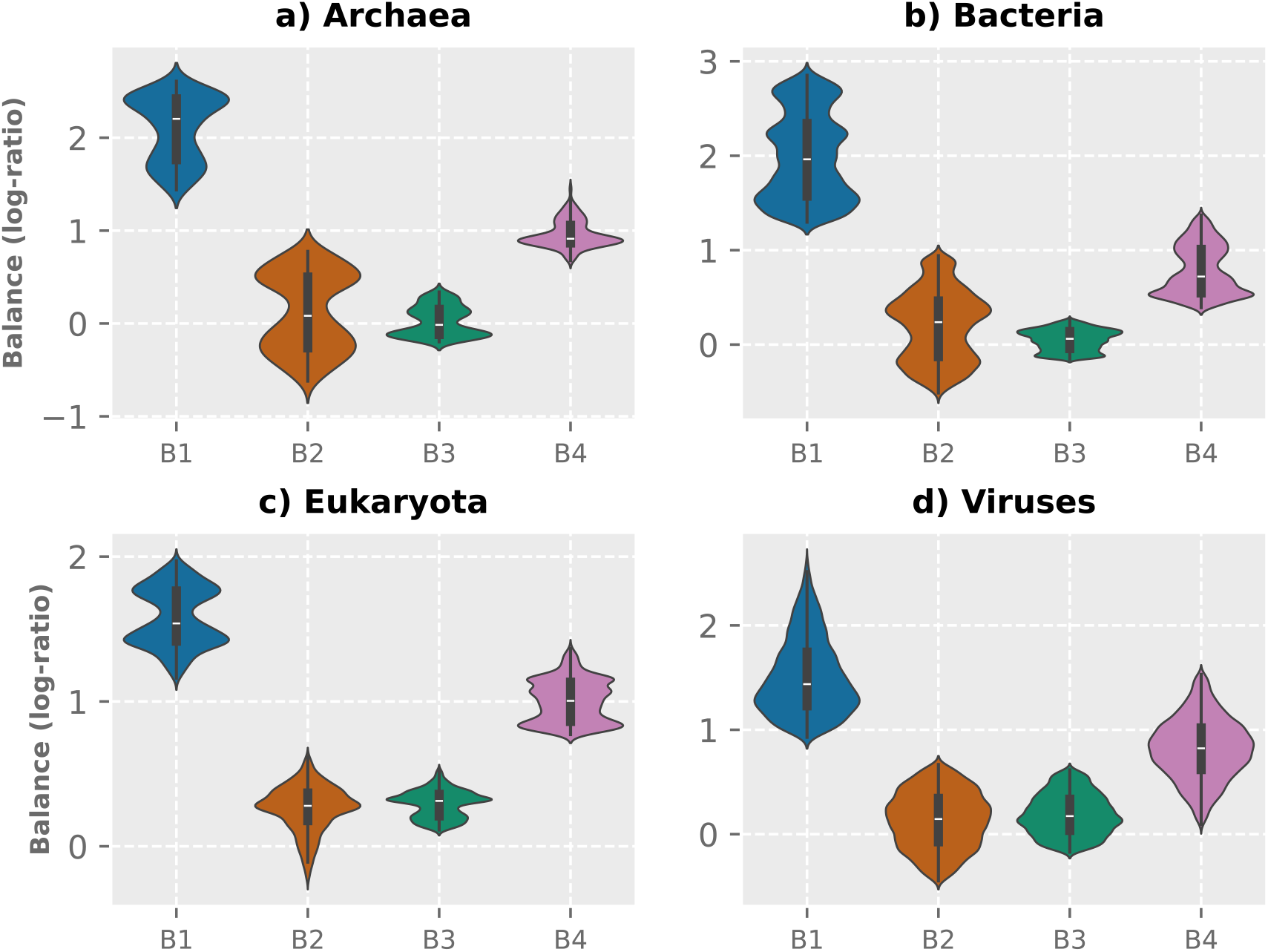
Distribution of balance values in protein-coding genes, aggregated by species. Blue, B1; orange, B2; green, B3; pink, B4. Balances below the 5^th^ and above the 95^th^ percentile are excluded to improve visualization.

### Assigned Numbers of Synonymous Codons Matches Genetic Codes

Finally, we calculated the optimal number of codons (by Integer Linear Programming, see Methods) for each amino acid based on the observed frequencies (Figure 9). Overall, the codon assignments generally agree with those based on genetic codes, with some notable deviations, as shown in Figure 4. Cysteine and histidine are assigned one codon, while aspartate and glutamate are assigned twice as many codons (up to five codons, depending on the domain) as they normally have in most codes. The same is true for alanine in archaea and bacteria, where a high abundance of this amino acid is observed. Of the three amino acids with high number of synonymous codons (leucine, arginine, and serine), only leucine is assigned six codons in bacteria and eukaryotes. Arginine and serine receive only about half as many codons in all domains (except for serine in eukaryotes). Interestingly, although methionine is generally overabundant in all domains, it is assigned only one codon (except in viruses), which is consistent with most genetic codes.

**Figure 9.**
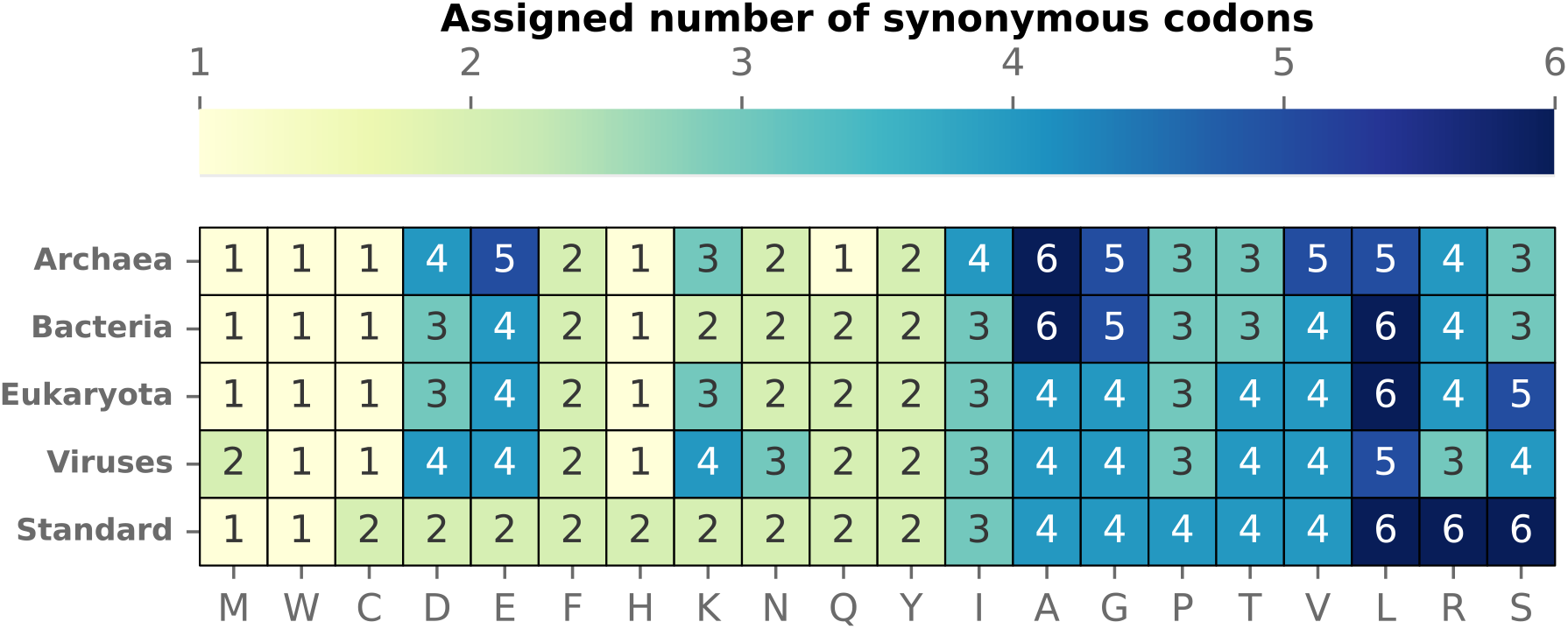
Optimal number of codons for each amino acid based on the observed frequencies (taken as the median over each domain) as well as given in the standard genetic code.

Looking at the overall median Jensen-Shannon divergence between amino acid frequencies based on the given genetic codes and predicted codon degeneracy distributions, the values for all domains are close to zero and generally decrease from archaea (codon: 0.032, GC: 0.029) to bacteria (codon: 0.024, GC: 0.026) and viruses (codon: 0.021, GC: 0.019) to eukaryotes (codon: 0.014, GC: 0.014) (Figure 10).

**Figure 10.**
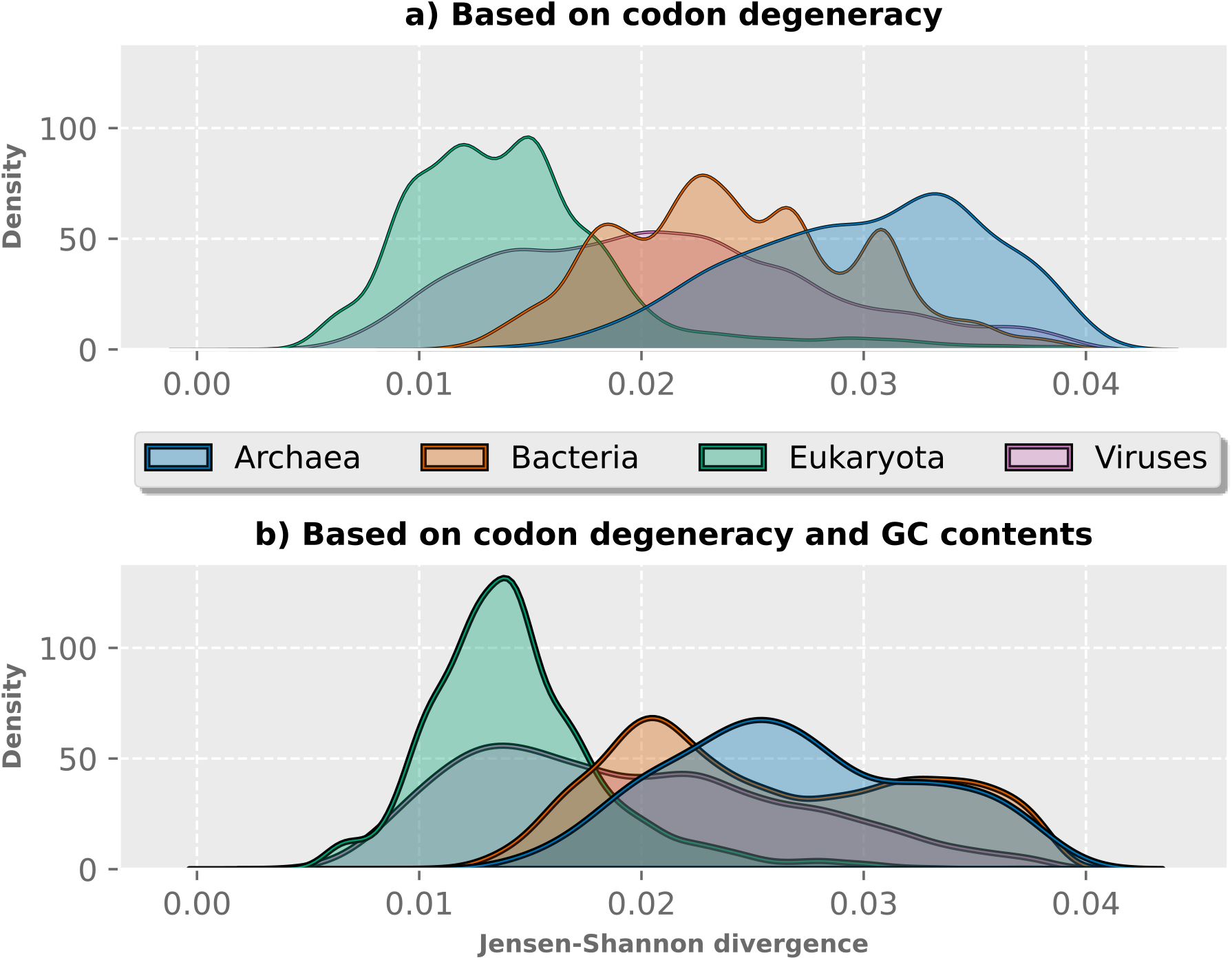
Jensen-Shannon divergences between observed and predicted frequencies using ILP for all 20 canonical amino acids. a) Based on codon degeneracy alone, b) based on codon degeneracy adjusted for GC content. Divergences above the 95^th^ percentile are excluded to improve visualization. Blue, archaea; orange, bacteria; green, eukaryotes; pink, viruses.

In the context of the constraint that the number of synonymous codons for each amino acid should be integers, the following question is of interest:

• How many of the 20 canonical amino acids can at most be encoded by *k* triplets in an alternative genetic code, assuming that at least one codon is needed for each amino acid and stop?

The solution can be found by the equation (*k* − 1) ∗ *x* + 20 = 64 − #*Stops*, which has the solution 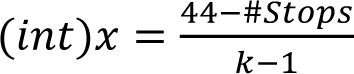, where *x* is the highest number of amino acids with at least *k* synonymous codons. For example, for *k* = 6 and three stop codons, this equation gives *x* = 8. Thus, not only leucine, serine and arginine could have six codons, but also another five amino acids, if all others have a single codon each and one has two.

## Discussion

The analysis of the relationships between number of synonymous codons, genomic composition, and amino acid occurrence in protein-coding genes offers valuable insights into the evolution of proteomes. To the best of our knowledge, the present study is the most comprehensive analysis of the correlation between codon degeneracy and amino acid frequency so far. For the first time, it encompasses all species available in the UniProt Knowledgebase, thereby covering all domains of life as well as viruses.

It is important to note that the codon multiplicity represent compositional data consisting of whole numbers. Therefore, no strong correlations with amino acid frequencies are to be expected if these vary too greatly. In the hypothetical case where, for example, five amino acids each occur ten times as frequently as any of the remaining amino acids, their numbers of synonymous codon cannot be ten or one respectively, as the sum is limited to 64 (minus the number of stop codons). According to our results, the greatest ratios between the median amino acid frequencies are found between alanine and cysteine in archaea and bacteria (Archaea: 16, bacteria: 14.1) and between leucine and cysteine in eukaryotes and viruses (eukaryotes: 7.9, viruses: 6.9). This suggests that constraining codon multiplicity to integer values slightly weakens the correlations, though the effect remains minor.

Our results may contribute to resolve the chicken-and-egg problem mentioned in the Introduction, namely whether codon degeneracy adapted to amino acid demand or vice versa. Positive correlations between amino acid frequencies and codon multiplicity may be interpreted such that the genetic code adapted to the requirements, at least those in the prebiotic era. However, a striking observation is that the correlation is particularly strong in eukaryotes although they evolved much later than the genetic code.

In view of our results, Francis Crick’s famous concept of the genetic code as a “frozen accident” can be revisited [11,21]. It does not seem to be coincidental but rather shaped by differences in the demand for specific amino acids. All deviations from the strong correlation are likely to have mainly evolved in the second era of evolution. Additionally, in that phase, optimization of amino acid frequencies has also been achieved by changes in the codon bias.

We have analyzed the amino acid frequencies without any prior assumptions such as codon bias or complexity of proteins. If all codons (except stop codons) were distributed uniformly, the strong correlation between codon degeneracy and amino acid frequency would result naturally. However, that distribution is not uniform due to codon bias and preferences for certain amino acids [64,65]. Interestingly, it can be concluded from our results that codon bias does not significantly affect the distribution of amino acids.

Several structural proteins show a low complexity. For example, collagen is mainly composed of proline and glycine [66,67], and silk fibroin is mainly composed of glycine, alanine and serine [68]. Our results indicate that biases of this kind average out over the entire genome-derived proteome, which is partly due to the fact that structural proteins constitute a small minority therein. Most proteins (such as enzymes, regulatory proteins, receptors etc.) have a much higher sequence complexity.

Our computational analysis based on data mining disregards gene expression so that all genes enter the analysis only in proportion to their length but not to their expression level. Furthermore, we neglect post-translational modification.

While our results are largely in line with our working hypothesis that amino acid frequency reflects the number of synonymous codons, significant deviations reveal additional influences, such as biochemical, structural, and ecological factors, shaping proteome composition. In particular, consistent deviations in eight amino acids across the four domains deserve further exploration of their underlying causes. Cysteine, arginine, histidine, and proline are consistently underrepresented in view of their codon degeneracy, while glutamate, aspartate, lysine, and methionine show persistent overrepresentation. Interestingly, among the amino acids with notable deviations, five are charged, as has been found earlier for specific organisms [6,7]. This may be related to the necessity to keep cytosolic proteins in solution and the properties of catalytic centres of enzymes. Adjusting for GC content reduces some deviations, especially in viruses, indicating that genomic composition significantly affects amino acid frequencies [26].

The consistency of these patterns across domains suggests universal evolutionary pressures influencing amino acid composition beyond codon degeneracy alone. When it comes to (deviations in) codon frequency and its impact on the proteome, Jacques Monod’s famous phrase that “What is true for *E. coli*, is true for the elephant” largely applies [69].

Proline, which is underrepresented, imposes structural rigidity that can constrain protein folding and dynamics, thereby limiting its incorporation in diverse contexts [70,71]. It is worth noting that in our study, we only analyzed the occurrence of amino acids in proteins. Moreover, they often occur in free form, for example, proline as an osmoprotectant in plants or glutamate as a neurotransmitter in animals [72,73].

Cysteine, which is strikingly underrepresented across all domains, requires sulphur – a nutrient often scarce in the environment [4,74]. Furthermore, that cysteines spontaneously form sulphur bridges under oxidative conditions, which may cause protein misfolding, aggregation, and loss of function if they are too abundant. This could explain the difference in abundance between cysteine and methionine, which involves sulphur as well. The overrepresentation of methionine may stem from its essential role as the translation initiator [74,75]. Note that we did not consider the excision of the initial methionine in many proteins, which allows cells to recycle this costly amino acid.

As mentioned above, alanine is the most abundant amino acid in archaea and bacteria and remains highly prevalent across other domains. It serves primarily as a structural component that promotes backbone stability in proteins, with few specific functions aside from potential interactions with lipophilic ligands. Although glycine is metabolically inexpensive to synthesize, it is less abundant than alanine, likely due to its non-chirality and its short side chain, which disrupts α-helix formation [75]. However, glycine is frequently found in β-sheets, such as those in β-keratin [76].

Interestingly, the magnitude of deviations varies among domains. For example, cysteine is more underrepresented in archaea and bacteria than in eukaryotes and viruses. Viruses, which rely on the translation machinery of the host, exhibit patterns influenced by host codon usage preferences, though some deviations persist, indicating intrinsic viral constraints.

The evolutionary timeline of amino acid incorporation into the genetic code offers additional context [17,77]. The temporal sequence of amino acid recruitment is significant. Amino acids acquired early in evolution, such as glycine, alanine, and leucine, had a broader selection of codons, whereas later-recruited amino acids, including methionine, cysteine, and tryptophan, had fewer remaining codons be assigned to them. The observed abundances of glutamate, lysine, and aspartate, despite their encoding by only two codons, are likely to be caused by additional pressures beyond codon availability, such as metabolic demand and protein functionality. By balancing the number of synonymous codons for each amino acid with their metabolic and functional demands, the genetic codes ensure the adaptability of proteomes across diverse organisms.

Krick and co-workers proposed that, in analogy to thermodynamics, a formal free energy composed of energetic synthesis costs and entropy of protein sequence variability was minimized during evolution [5]. An illustrative example is silk fibroin, which is mainly composed of cheap amino acids (see above). In that case, the energetic parameter dominates. In contrast, in enzymes and regulatory proteins, another compromise between these two factors was found, in which the entropic parameter has a considerable influence. The correlation to synthesis costs is certainly worth investigating further. Other factors include selection for protein properties like charge balance and flexibility, post-translational modifications, and evolutionary drift or mutation biases.

The charge balance need not be perfect because proteins (in contrast to the entire cell) need not be electroneutral and are compensated by counter-ions such as chloride and potassium [78]. Here, we have analyzed both the total frequency of positively charged and that of the negatively charged amino acids at neutral pH (Figure 7). It is worth noting, though, that the pH values depend on cell type, intra- or extracellular localization, external factors etc. Moreover, as mentioned above, expression levels are ignored here.

To sum up, there is a strong correlation between the number of synonymous codons and amino acid frequency across domains, with some notable deviations, which deserve further studies. These may be related to the constraint on the number of synonymous codons to be integers and to codon usage bias. The latter introduces additional complexity, as organisms optimize for tRNA availability, favouring more efficiently translated codons and, consequently, specific amino acids [79,80]. Horizontal gene transfer introduces variability by inheriting codon usage patterns from donors [81]. Environmental pressures in habitats shape amino acid usage to enhance protein stability [19].

Moreover, when considering the influence of the GC content, we have assumed that G and C occur equally in the single strand and so do A and T, as expressed in Chargaff’s second parity rule [42]. Our analysis can be extended to cases where that rule does not hold, such as in mitochondria, plastids and some viruses. Furthermore, our results may be interesting in view of synthetic biology [82]. When designing synthetic genomes, a relevant question is whether the correlation under study should be maintained or whether deviating from it offers advantages.

## Author Contribution Statement

V.W, G.T, and S.S. designed the study. V.W. performed the data analysis and created all figures and tables. V.W. and S.S. drafted the manuscript. All authors contributed to the interpretation of the results and revised the manuscript.

## Supporting information

All figures as PDF

All tables as DOCX

## Acknowledgments

We thank Jan Grau (Halle) and Steve Hoffmann (Jena) for stimulating discussions and Gabriel Lencioni Lovate (Jena) for helpful comments on the manuscript. Financial support by the German Research Foundation (DFG) in the SFB 1127 ChemBioSys project no. 239748522 to S.S. is gratefully acknowledged. During the preparation of this work, the authors used Google Gemini 3.6 Flash and GPT-5.6 for text optimisation.

## Availability of Data and Materials

The datasets generated and analysed during this study along with the code and several supplementary files are available in the Mendeley repository: Wesp, Valentin; Theißen, Günter; Schuster, Stefan (2026), “Strong correlation between amino acid frequency and codon degeneracy in genetic codes across all domains of life”, Mendeley Data, V2, doi: 10.17632/w5nycn4cdx.2

