## Supplementary figures and images for "Strong correlation between amino acid frequency and codon degeneracy in genetic codes across all domains of life"

### Figure 1.pdf

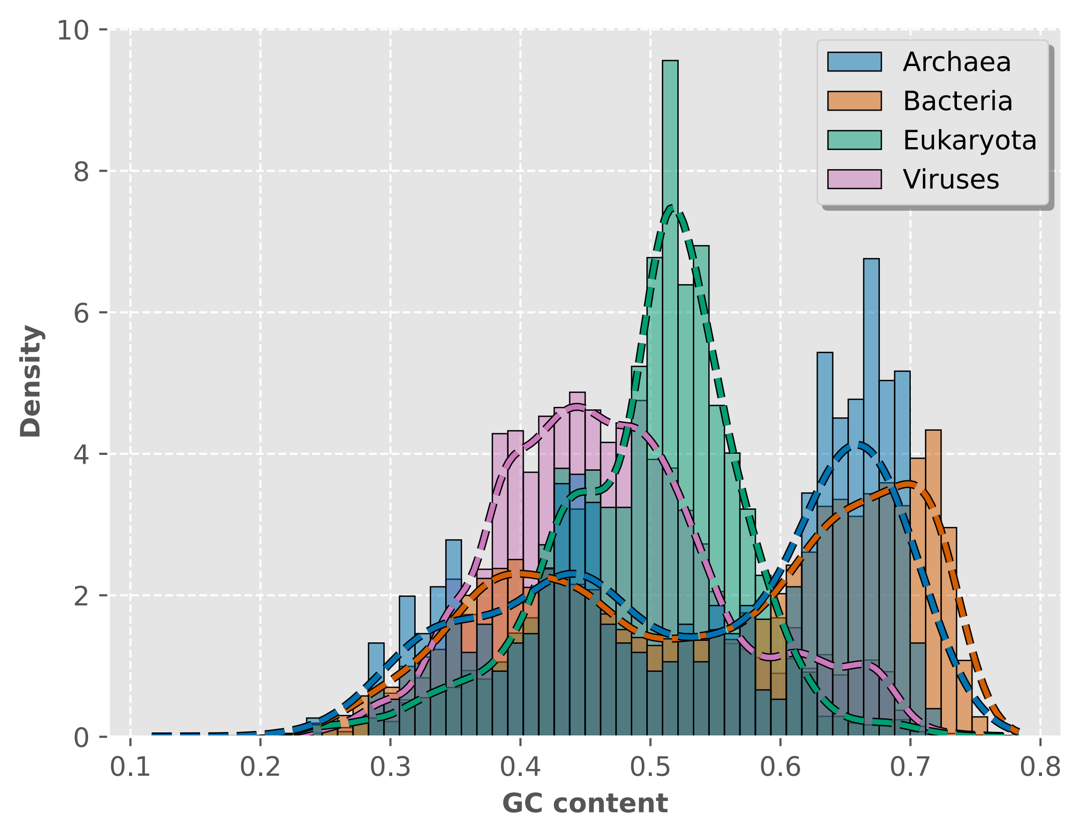

### Figure 2.pdf

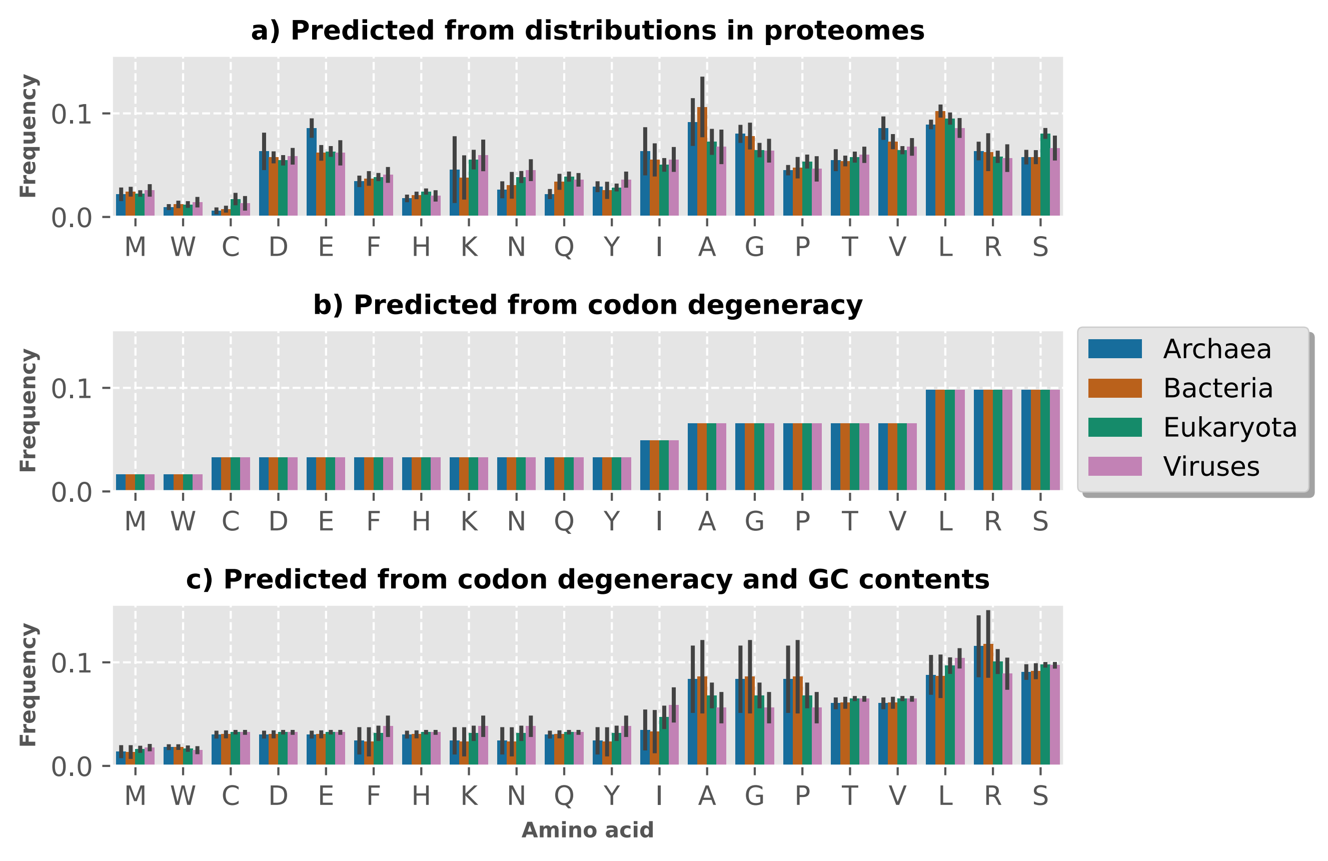

### Figure 3.pdf

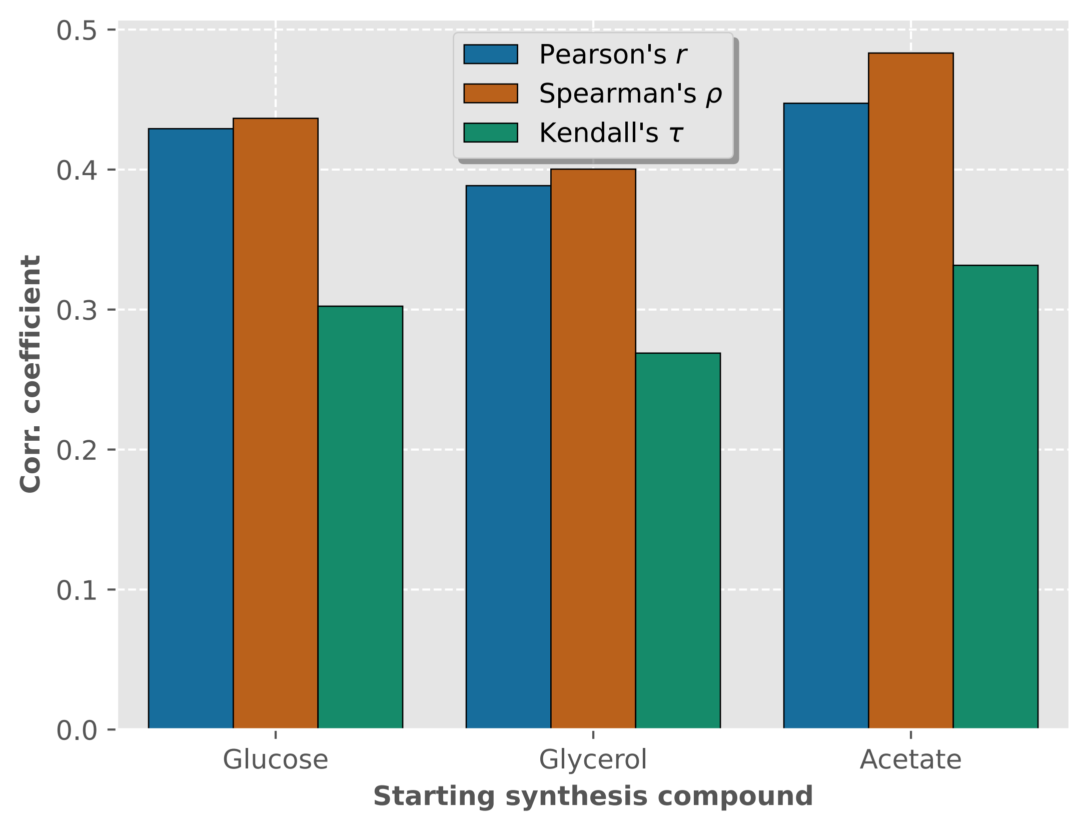

### Figure 4.pdf

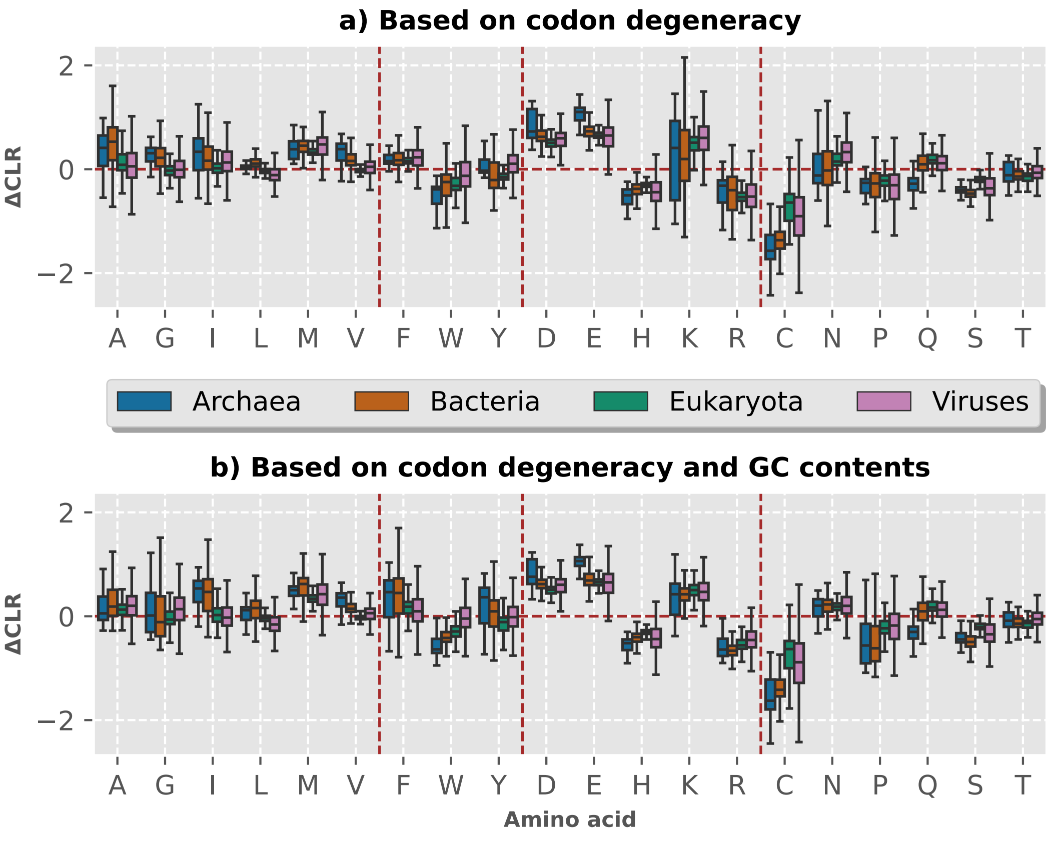

### Figure 5.pdf

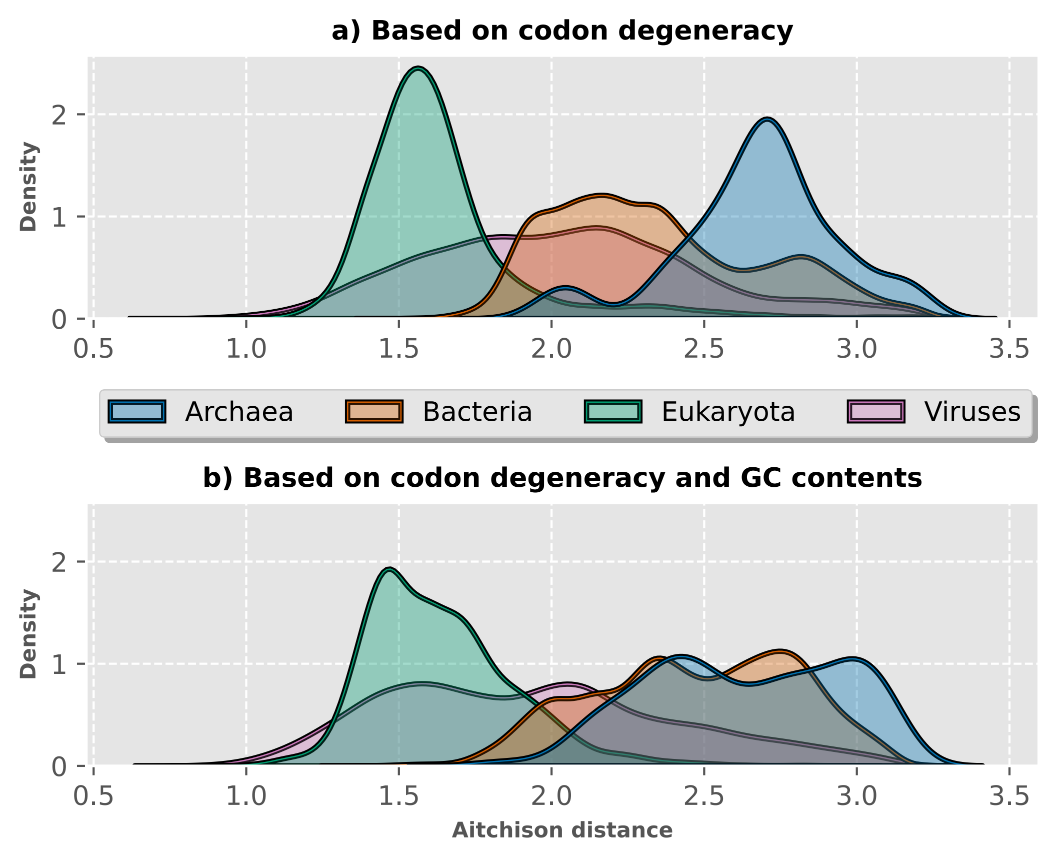

### Figure 6.pdf

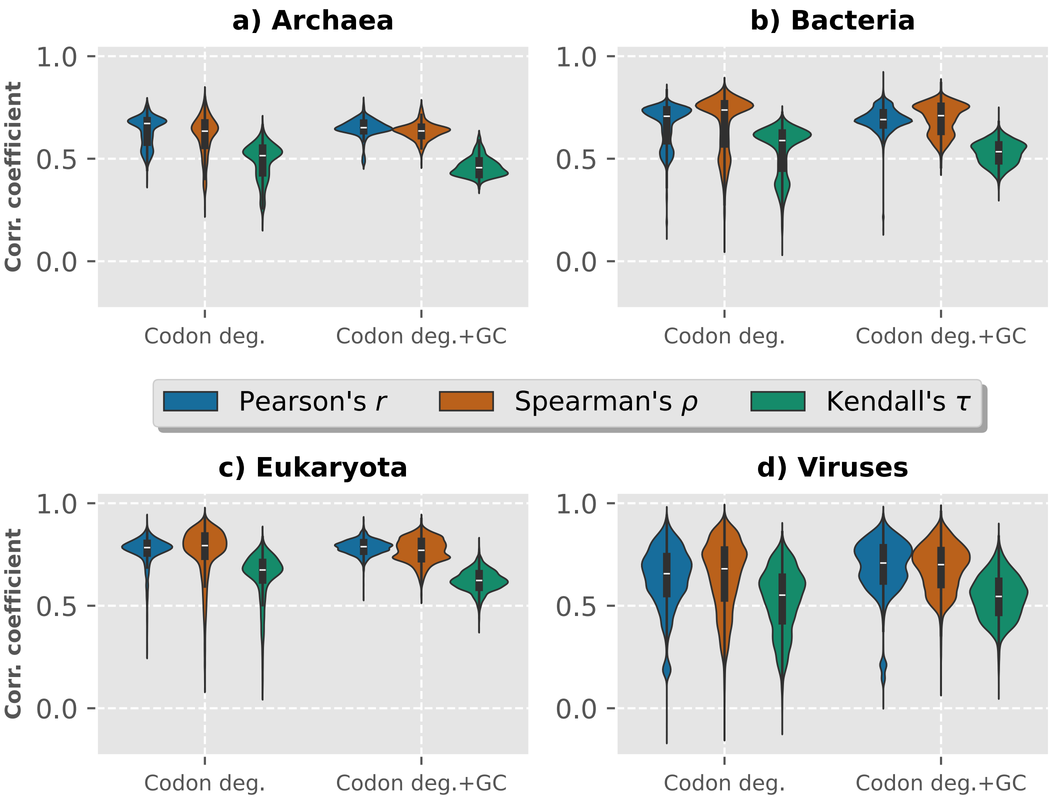

### Figure 7.pdf

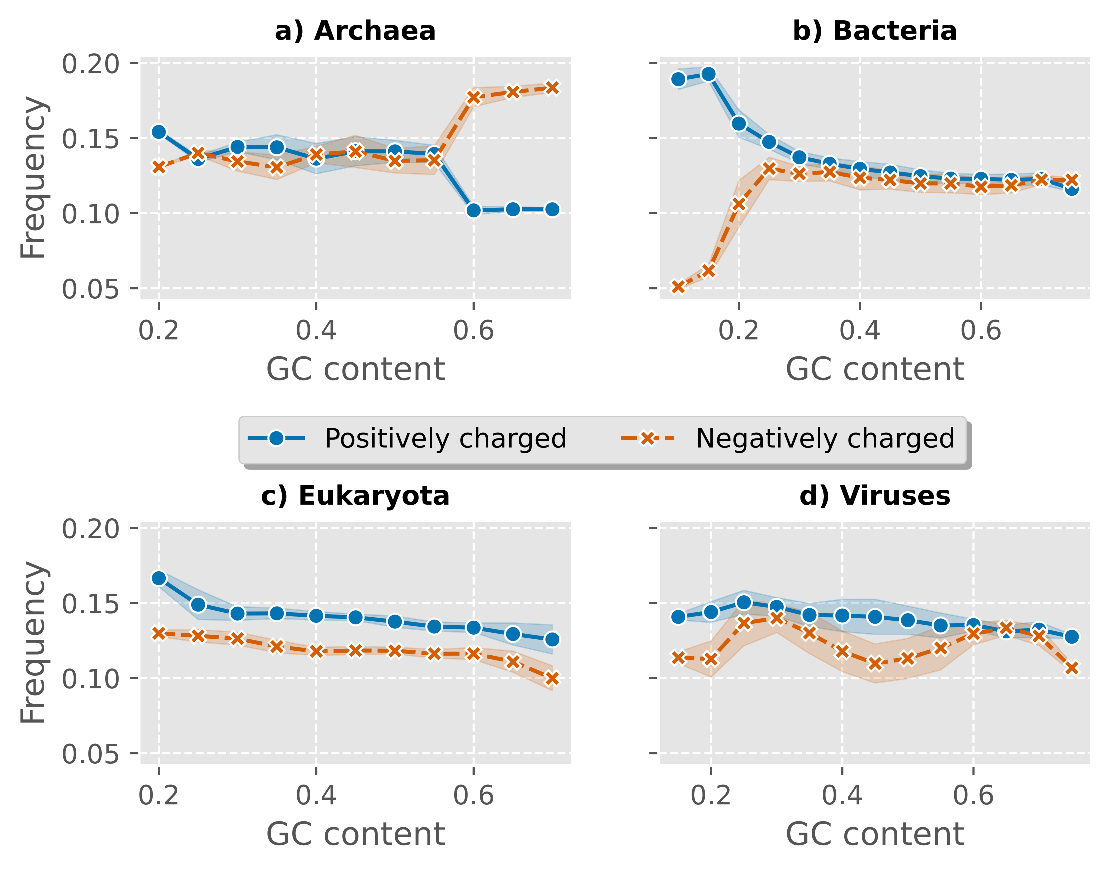

### Figure 8.pdf

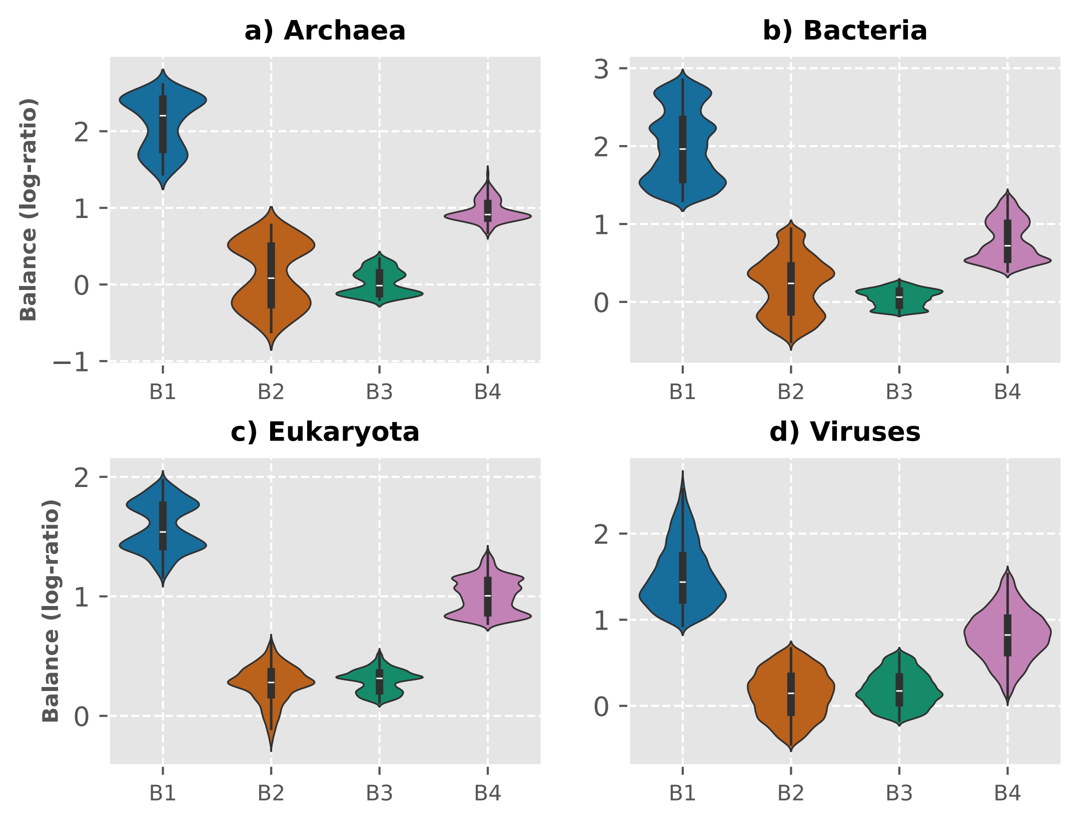

### Figure 9.pdf

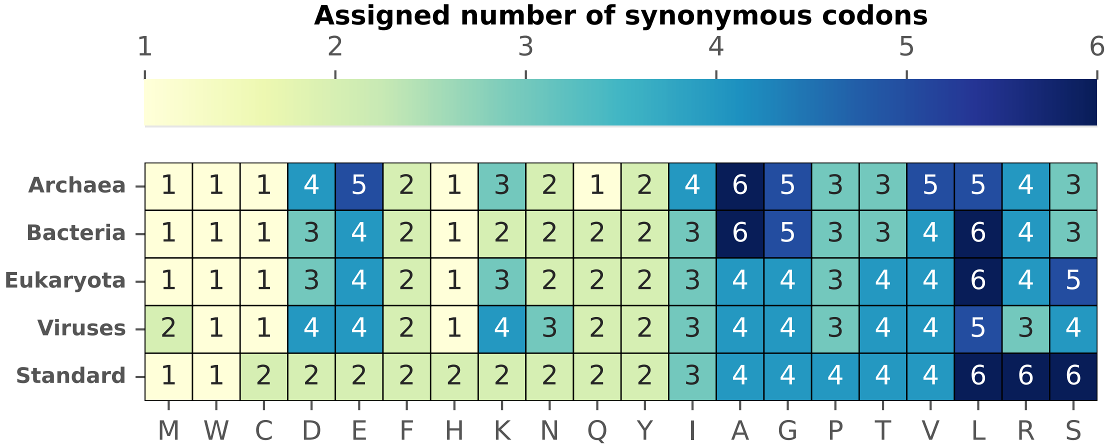

### Figure 10.pdf

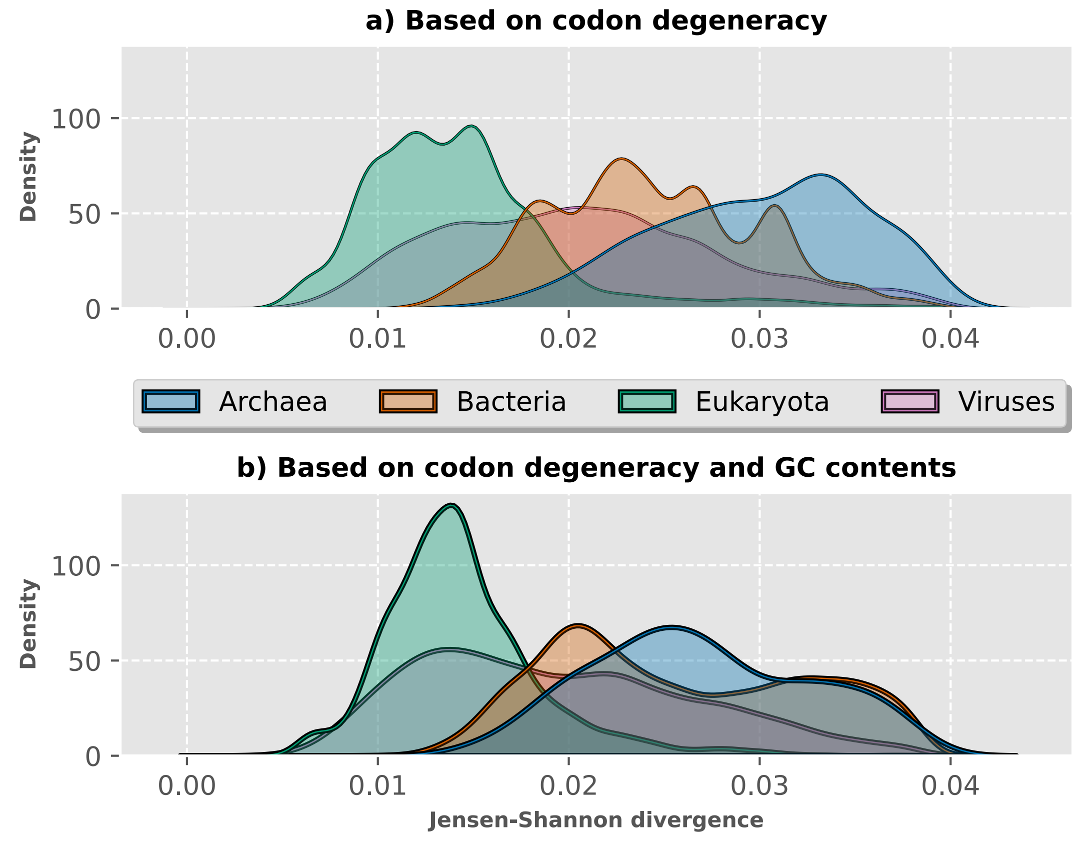

### Figure S1.pdf

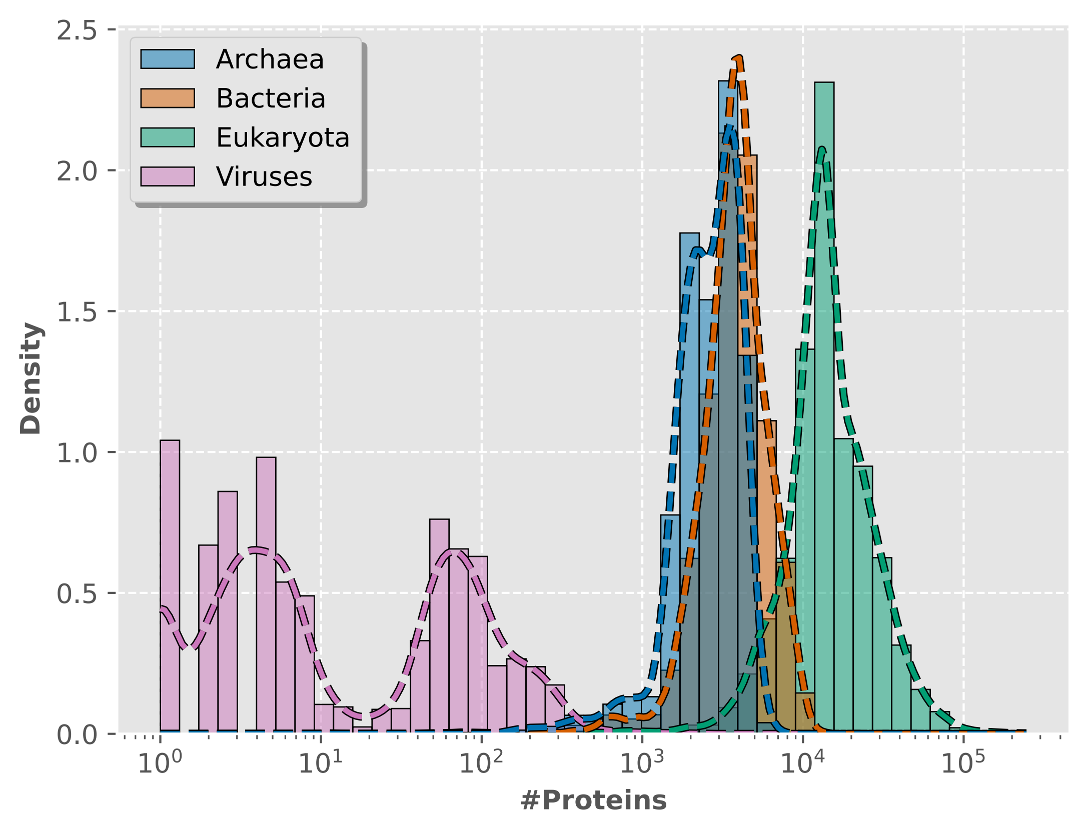

### Figure S2.pdf

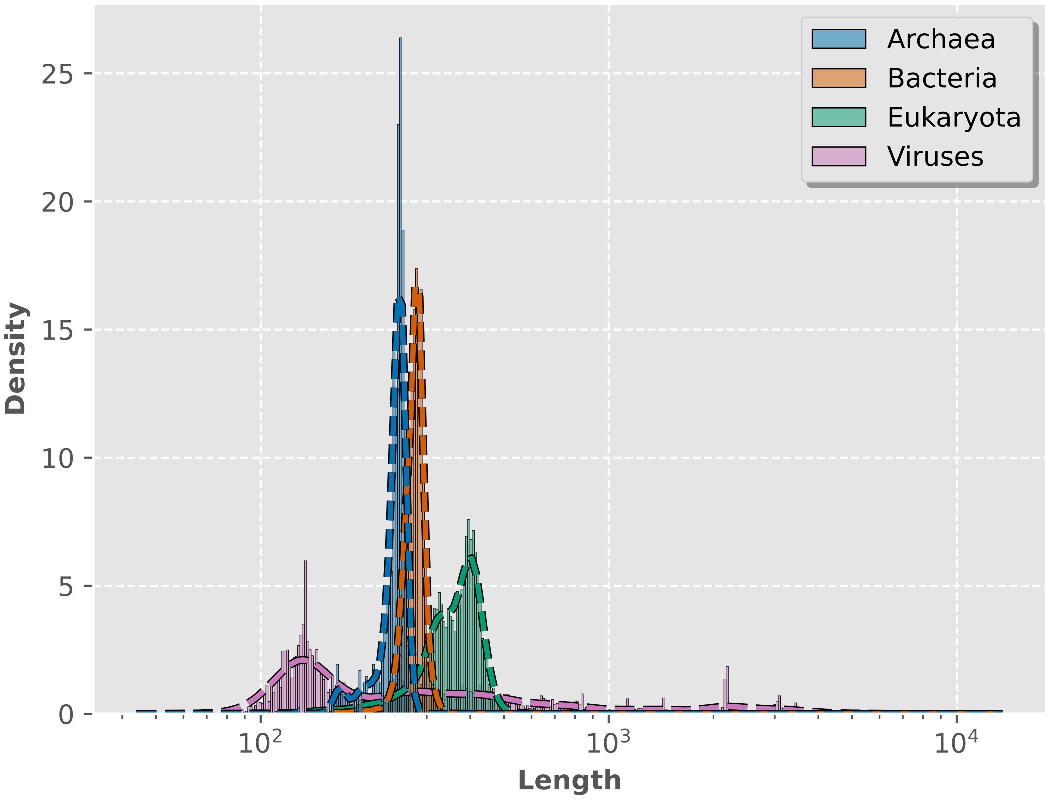

### Figure S3.pdf

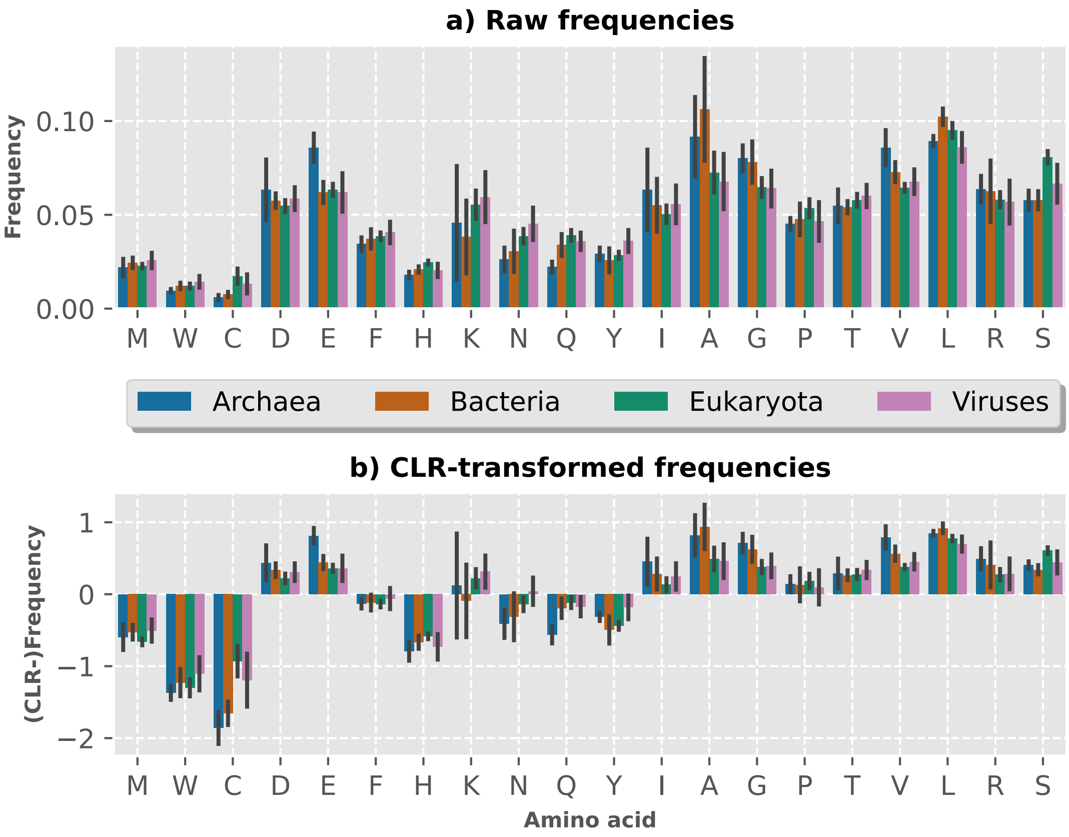

### Figure S4.pdf

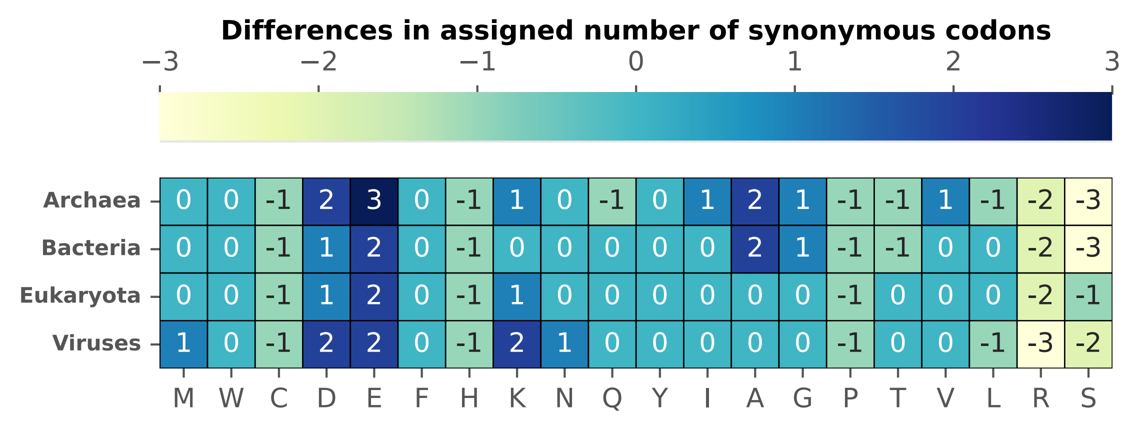
