## Supplementary material for "Strong correlation between amino acid frequency and codon degeneracy in genetic codes across all domains of life": All tables as DOCX: Table 1.docx

**Table 1.** Distribution of proteomes and proteins (count and median) for each domain as available from the UniProt Knowledgebase [39].

| **Domain** | **#Proteomes** | **#Proteins** | **x̃Proteins** |
| --- | --- | --- | --- |
| **Archaea** | 634 | 1,754,750 | 2,710 |
| **Bacteria** | 17,903 | 73,458,183 | 3,812 |
| **Eukaryotes** | 3,498 | 61,332,190 | 13,882.5 |
| **Viruses** | 14,335 | 681,211 | 7 |
