## Supplementary material for "Strong correlation between amino acid frequency and codon degeneracy in genetic codes across all domains of life": All tables as DOCX: Table 2.docx

**Table 2**. Distribution of genetic codes extracted for each domain from the NCBI taxonomy browser [40,41], with IDs sourced from the NCBI genetic code database [35].

| **Code ID** | **Archaea** | **Bacteria** | **Eukaryotes** | **Viruses** |
| --- | --- | --- | --- | --- |
| **1** | 0 | 0 | 3,431 | 7,632 |
| **4** | 0 | 182 | 1 | 100 |
| **5** | 0 | 0 | 0 | 0 |
| **6** | 0 | 0 | 15 | 2 |
| **10** | 0 | 0 | 2 | 0 |
| **11** | 634 | 17,718 | 0 | 6,571 |
| **12** | 0 | 0 | 48 | 0 |
| **15** | 0 | 0 | 0 | 8 |
| **16** | 0 | 0 | 0 | 21 |
| **25** | 0 | 1 | 0 | 0 |
| **26** | 0 | 0 | 1 | 0 |
