## Supplementary material for "Strong correlation between amino acid frequency and codon degeneracy in genetic codes across all domains of life": All tables as DOCX: Table 3.docx

**Table 3**. The recursive splits for the calculations of balances (**B1** to **B4**) based on the different number of codons in the standard genetic code^*^.

| **B1** | **B2** | **B3** | 6 |
| --- | --- | --- | --- |
|  |  |  | 4 |
|  |  |  | 3 |
|  | **B4** | | 2 |
|  |  |  | 1 |

*^*^*In each $B_{i}$ comparison, the log-ratios of the two subsets to the right of $B_{i}$ are compared with one another.
