## Supplementary material for "Strong correlation between amino acid frequency and codon degeneracy in genetic codes across all domains of life": All tables as DOCX: Table 4.docx

**Table 4.** Observed median amino acid distributions and their median absolute deviations (in %) in genome-encoded proteome across domains.

| **Amino acid** | **Archaea** | **Bacteria** | **Eukaryotes** | **Viruses** |
| --- | --- | --- | --- | --- |
| **M** | 2.19 (±0.5) | 2.44 (±0.3) | 2.27 (±0.1) | 2.56 (±0.4) |
| **W** | 0.95 (±0.1) | 1.21 (±0.2) | 1.2 (±0.1) | 1.43 (±0.3) |
| **C** | 0.59 (±0.1) | 0.75 (±0.1) | 1.73 (±0.4) | 1.32 (±0.5) |
| **D** | 6.33 (±1.6) | 5.76 (±0.4) | 5.47 (±0.3) | 5.86 (±0.6) |
| **E** | 8.58 (±0.7) | 6.19 (±0.6) | 6.33 (±0.3) | 6.19 (±1) |
| **F** | 3.44 (±0.4) | 3.72 (±0.5) | 3.86 (±0.2) | 4.06 (±0.6) |
| **H** | 1.79 (±0.2) | 2.08 (±0.2) | 2.46 (±0.1) | 2.04 (±0.4) |
| **K** | 4.57 (±3) | 3.82 (±1.9) | 5.54 (±0.8) | 5.94 (±1.3) |
| **N** | 2.62 (±0.6) | 3.06 (±1.1) | 3.86 (±0.4) | 4.52 (±0.9) |
| **Q** | 2.21 (±0.3) | 3.39 (±0.6) | 3.9 (±0.3) | 3.59 (±0.5) |
| **Y** | 2.92 (±0.3) | 2.57 (±0.6) | 2.85 (±0.2) | 3.6 (±0.6) |
| **I** | 6.34 (±2.1) | 5.51 (±1.4) | 5.03 (±0.5) | 5.56 (±1) |
| **A** | 9.17 (±2.1) | 10.62 (±2.7) | 7.25 (±1.1) | 6.77 (±1.5) |
| **G** | 8.02 (±0.7) | 7.81 (±1.1) | 6.46 (±0.5) | 6.4 (±1) |
| **P** | 4.52 (±0.3) | 4.76 (±0.8) | 5.35 (±0.5) | 4.64 (±1) |
| **T** | 5.49 (±0.9) | 5.41 (±0.3) | 5.78 (±0.3) | 6 (±0.6) |
| **V** | 8.58 (±0.9) | 7.27 (±0.5) | 6.45 (±0.2) | 6.76 (±0.7) |
| **L** | 8.93 (±0.3) | 10.23 (±0.4) | 9.5 (±0.4) | 8.59 (±0.8) |
| **R** | 6.37 (±0.7) | 6.26 (±1.6) | 5.81 (±0.4) | 5.68 (±1.1) |
| **S** | 5.78 (±0.5) | 5.78 (±0.5) | 8.07 (±0.3) | 6.65 (±1) |
