## Supplementary material for "Strong correlation between amino acid frequency and codon degeneracy in genetic codes across all domains of life": All tables as DOCX: Table 5.docx

**Table 5.** Biosynthesis yields (mol/mol) for each amino acid (LP Std) in E. coli (taxonomy ID: 83333) from glucose, glycerol, and acetate [33] as well as median amino acid distribution as determined in our analysis (in %).

| **Amino acid** | **Glucose** | **Glycerol** | **Acetate** | **Observed**  **abundance** |
| --- | --- | --- | --- | --- |
| **M** | 0.62 | 0.36 | 0.16 | 2.91 |
| **W** | 0.47 | 0.25 | 0.12 | 1.34 |
| **C** | 1.03 | 0.6 | 0.27 | 1.11 |
| **D** | 1.86 | 1 | 0.5 | 5.3 |
| **E** | 1.15 | 0.6 | 0.33 | 6.08 |
| **F** | 0.56 | 0.3 | 0.14 | 3.75 |
| **H** | 0.89 | 0.49 | 0.22 | 2.23 |
| **K** | 0.8 | 0.47 | 0.21 | 4.45 |
| **N** | 1.74 | 1 | 0.47 | 3.81 |
| **Q** | 1.19 | 0.61 | 0.33 | 4.35 |
| **Y** | 0.58 | 0.3 | 0.15 | 2.69 |
| **I** | 0.75 | 0.44 | 0.2 | 6.07 |
| **A** | 2 | 1 | 0.5 | 9.54 |
| **G** | 2.73 | 1.54 | 0.71 | 7.25 |
| **P** | 1.01 | 0.57 | 0.29 | 4.36 |
| **T** | 1.3 | 0.75 | 0.35 | 5.36 |
| **V** | 1 | 0.5 | 0.25 | 7.23 |
| **L** | 0.75 | 0.38 | 0.22 | 10.75 |
| **R** | 0.89 | 0.51 | 0.25 | 5.65 |
| **S** | 2 | 1 | 0.5 | 5.78 |
