## Supplementary material for "Strong correlation between amino acid frequency and codon degeneracy in genetic codes across all domains of life": All tables as DOCX: Table 6.docx

**Table 6.** Median deviations between empirical values and theoretical values derived from codon degeneracy and from codon degeneracy adjusted for GC content for the 20 canonical amino acids in CLR space.

|  | **Archaea** | | **Bacteria** | | **Eukaryotes** | | **Viruses** | |
| --- | --- | --- | --- | --- | --- | --- | --- | --- |
| **Amino acid** | **Code** | **GC** | **Code** | **GC** | **Code** | **GC** | **Code** | **GC** |
| **A** | 0.41 | 0.05 | 0.53 | 0.19 | 0.09 | 0.13 | 0.06 | 0.2 |
| **G** | 0.31 | 0.01 | 0.22 | -0.11 | -0.03 | -0.06 | -0.01 | 0.14 |
| **I** | 0.34 | 0.53 | 0.16 | 0.47 | 0.02 | 0.02 | 0.13 | -0.03 |
| **L** | 0.04 | 0.12 | 0.11 | 0.16 | -0.04 | -0.04 | -0.11 | -0.16 |
| **M** | 0.38 | 0.5 | 0.45 | 0.62 | 0.32 | 0.34 | 0.48 | 0.43 |
| **V** | 0.39 | 0.36 | 0.16 | 0.14 | -0.03 | -0.03 | 0.05 | 0.06 |
| **F** | 0.16 | 0.46 | 0.17 | 0.45 | 0.15 | 0.18 | 0.23 | 0.1 |
| **W** | -0.39 | -0.64 | -0.25 | -0.42 | -0.32 | -0.29 | -0.13 | -0.04 |
| **Y** | -0.03 | 0.36 | -0.21 | 0.09 | -0.15 | -0.11 | 0.11 | -0.02 |
| **D** | 0.73 | 0.76 | 0.62 | 0.62 | 0.51 | 0.51 | 0.6 | 0.61 |
| **E** | 1.1 | 1.06 | 0.73 | 0.69 | 0.65 | 0.65 | 0.65 | 0.65 |
| **H** | -0.5 | -0.52 | -0.38 | -0.4 | -0.3 | -0.3 | -0.44 | -0.44 |
| **K** | 0.41 | 0.42 | 0.2 | 0.42 | 0.51 | 0.5 | 0.6 | 0.47 |
| **R** | -0.32 | -0.64 | -0.4 | -0.66 | -0.54 | -0.56 | -0.53 | -0.46 |
| **C** | -1.57 | -1.63 | -1.37 | -1.42 | -0.64 | -0.63 | -0.9 | -0.89 |
| **N** | -0.12 | 0.2 | -0.02 | 0.23 | 0.15 | 0.19 | 0.33 | 0.2 |
| **P** | -0.26 | -0.56 | -0.27 | -0.62 | -0.22 | -0.23 | -0.31 | -0.18 |
| **Q** | -0.28 | -0.3 | 0.09 | 0.09 | 0.17 | 0.17 | 0.12 | 0.12 |
| **S** | -0.4 | -0.45 | -0.47 | -0.49 | -0.2 | -0.21 | -0.37 | -0.35 |
| **T** | -0.12 | -0.09 | -0.14 | -0.14 | -0.13 | -0.13 | -0.07 | -0.05 |
